# Early Ontogenetic Development of Tessellated Cartilage in Chondrichthyans

**DOI:** 10.64898/2026.08.21.746214

**Authors:** Hannah.M. Byrne, Isabel Breet, Bertie Joan van Heuven, Richard P. Dearden, Sophie Sanchez, Zerina Johanson, Mason N. Dean, Martin Rücklin

## Abstract

Tessellated cartilage (TC) is a hallmark of the chondrichthyan skeleton, yet its development early in ontogeny across the four major groups (batoids, galeomorphs, squalomorphs, and holocephalans) remains poorly understood. In this study, we used micro-CT scanning to examine TC in the jaws of ten representative taxa, allowing interspecific comparison and to determine whether specialised features previously observed in mature forms had yet developed Batoids consistently exhibited well-developed, polygonal tesserae regardless of feeding mode. This is in contrast to the heterogeneity in TC morphology of the galeomorphs. We report the first internal trabecular network in a non-batoid elasmobranch, *Ginglymostoma cirratum*, likely linked to mechanical pressures of suction feeding. *Mustelus mustelus* exhibited poor TC development despite being a durophagous feeder, pointing to a pronounced ontogenetic lag. *Squatina oculata* possessed significantly larger and thicker tesserae than other taxa in this study, likely related to its explosive ambush predation mode, whereas *Chimaera monstrosa* exhibited a poorly mineralised, mesh-like TC lacking discrete tesserae, consistent with previous findings Across all specimens, multi-layered TC was absent, confirming that multi-layering develops later in ontogeny. These results demonstrate that generalised models of TC development based on a few taxa is not representative of the heterogeneity in TC morphology.

## Introduction

Sharks, rays and their relatives (chondrichthyans) have roamed our oceans for over 430 million years and have survived four major mass extinctions (Bazzi et al., 2021; Friedman and Sallan, 2012; Guinot et al., 2013; Guinot and Condamine, 2023; Sallan and Coates, 2010). Extant species show remarkable morphological and ecological diversity, occupying habitats ranging from coastal shallows to pelagic depths beyond 3,700 metres and serve a significant ecological role in marine ecosystems as top-level predators (Cortés, 1999; Ebert, 2015; Hinrichs et al., 2026). This evolutionary success has been linked, in part, to a key innovation: a lightweight yet strong internal skeleton made predominantly of cartilage rather than bone(Kemp and Westrin, 1979). Palaeontological evidence, however, indicates that endochondral bone was present in the last common ancestors to chondrichthyans and their sister group, osteichthyans (bony fishes) arguing that chondrichthyans evolved their distinct and diverse cartilaginous tissues to deal with the functional skeletal demands met by bone in nearly all other vertebrate species (Brazeau et al., 2020).

The chondrichthyan skeleton is a composite of mineralised and unmineralised skeletal tissues. Whereas the vertebral column comprises densely mineralised tissues (Berio et al., 2021), the rest of the skeleton is predominantly an unmineralised hyaline-like cartilage, forming the bulk of most skeletal elements (Seidel et al., 2020b). A comparatively thin layer of calcified blocks known as **tesserae** forms a mosaic over the uncalcified cartilage core, with tesserae linked by collagen fibers and the tesserae mat overlain by a fibrous matrix called the perichondrium (Kemp and Westrin, 1979; Seidel et al., 2017). The layering of mineralised and unmineralised tissues gives the composite tissue its name: **Tessellated Cartilage** (TC) (Dean and Summers, 2006; Seidel et al 2016, 2020; Maisey et al., 2020). Tesserae blocks feature wedges of high mineral density which propagate from the centre of the tessera towards the joints with neighbouring tesserae, which are termed **spokes** (Jayasankar et al., 2020; Maisey et al., 2020a; Seidel et al., 2016). The spaces between neighbouring tesserae and spokes are termed **pores**. This is not to be confused with enlarged openings in the tesseral layer which are associated with mineralised struts traversing the uncalcified cartilage core to unite opposing tesseral layers, named trabeculae (Clark et al., 2022; Dean et al., 2006; Summers, 2000). These have also been referred to as pores in the literature (Clark et al., 2022). Because these larger openings and the spaces between tesserae have both been referred to as pores, we use specific terms to avoid confusion: **tesseral pores** for the spaces between tesserae and circumscribed by the spokes, and **canal pores** for the enlarged openings associated with trabeculae (see Figure 2 for examples). While the differences between the two have not been extensively explored, canal pores are typically larger, more circular, and explicitly linked to trabeculae (Clark et al., 2022; Jayasankar et al., 2020).

Shape and size of tesserae, canal and tesseral pores, and degree of mineralisation can differ significantly throughout the skeleton in both extant (Clark et al., 2022; Jayasankar et al., 2017; Seidel et al., 2020a, 2019; Wilga et al., 2024; Yang et al., 2024) and fossil taxa (Maisey, 2013; Maisey et al., 2020a; Pears et al., 2020), often in relation to skeletal element shape and/or habitat or dietary specialisations. For example, thickening of the tesseral layer is regularly associated with regions of high skeletal curvature and/or regions that experience high loads (e.g. beneath the teeth in species with hard prey diets; Clark et al 2022). While ‘normal’ tesserae are typically thin **polygonal** tiles, usually with 6 sides, this can vary to be 4-12 sides (Seidel et al., 2020b). In some species, fused tesserae have also been documented, where mineralisation of the fibrous perichondrium merges regions of tesserae into a single mass (Maisey et al., 2020; Clark et al., 2022), occasionally also pathologically (Dean et al., 2017; Seidel et al., 2017). Concurrently, regions of reduced tessellation have also been observed, where tesserae are either extremely thin and sparse (Yang et al., 2024) or reduced to just their connections to neighbouring tesserae. This results in **stellate** forms, termed trabecular tesserae by Atake et al. (2025) (see also Knötel et al., 2017; Jayasankar et al., 2020). Several specialised forms also exist, including enlarged, wedge-shaped **voussoir tesserae** or tall **columnar tesserae** found along ridges or foraminal margins and tend to lack tesseral pores (Clark et al., 2022; Maisey et al., 2020a). Tesserae can form a **multi-layered** crust on the skeleton’s cortex (Clark et al., 2022; Dean et al., 2017; Dingerkus et al., 1991). The early development of tessellated cartilage is nearly unexplored (Atake et al., 2025; Criswell et al., 2017; Seidel et al., 2016; Yang et al., 2024). As a result, it is currently unknown whether specialised tesseral features emerge early in ontogeny or are shaped only later in life (e.g. due to functional demands), although recent work suggests the tessellation pattern could be a function of the spatial distribution of initial growth centers in the tissue (Yang et al., 2024). Our study will investigate the early development of TC in a range of chondrichthyan taxa at neonate/juvenile stages to test these different possibilities.

Traditionally, calcified tissues in chondrichthyans have been studied using stained thin sections for light microscopy and scanning electron microscopy (Berio et al., 2021; Debiais-Thibaud, 2019; Kemp and Westrin, 1979; Pears et al., 2020; Schaefer and Summers, 2005; Summers, 2000), resulting primarily in two-dimensional data. Although these approaches have yielded important insights, they do not allow for observations of variation in calcified cartilage across entire skeletal elements, nor do they capture three-dimensional variation within the tesseral layer. In this regard, micro-computed tomography (micro-CT) has supported advances in the study of chondrichthyan skeletal morphology at resolutions adequate to resolve tesserae (e.g. Summers, 2000; Summers et al., 2004; Underwood et al., 2015, 2016), with new advances in digital image segmentation permitting the isolation and quantification of individual tesserae in histological detail (Chaumel et al., 2020; Clark et al., 2022; Seidel et al., 2020a, 2016). In this study, we characterise the tessellation of the palatoquadrate (herein referred to as the upper jaw, except in the holocephalan detailed below) and Meckel’s cartilage (i.e. lower jaw) from neonates/juveniles of taxa spanning the four major chondrichthyan groups: squalomorph and galeomorph sharks, batoids (rays and relatives), and holocephalans (chimaera) (Figure 1). This selection also encompasses a diversity of feeding modes. We visualise the tesseral layer in 3D in complete jaws using micro-CT scanning and 3D imaging, documenting the size, shape, and thickness of tesserae and presence of specialised features in a qualitative and quantitative manner. To our knowledge, this is the first comparison of tesseral variation over an entire skeletal element among several chondrichthyan taxa. This allows a unique opportunity to identify patterns and specialisations in tesserae morphology, and to test whether these can be explained by phylogenetic or mechanical constraints in jaw architecture, by feeding mode or by other factors.

**Figure 1.**
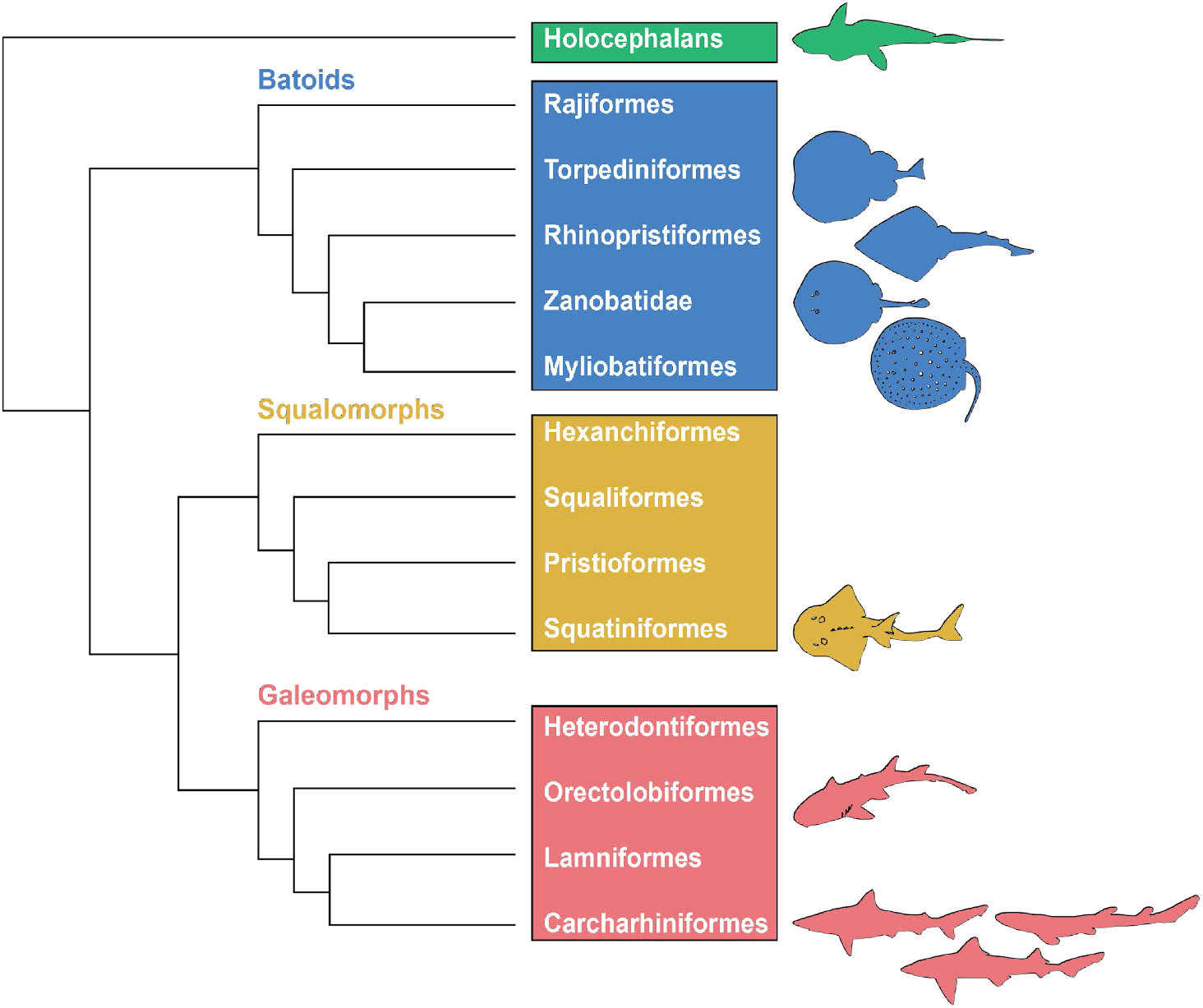
A simplified phylogeny of extant chondrichthyans (Aschliman et al., 2012; Capretz Batista da Silva et al., 2021; Naylor et al., 2012), with the four major superorders and their respective orders shown. Silhouettes represent the taxa studied and are obtained from PhyloPics.

## Materials and Methods

### Specimen selection

Specimens for this study were obtained from the wet collections at Naturalis Biodiversity Center, The Netherlands. In total, jaws from 12 chondrichthyan taxa were examined. These were selected to represent a broad phylogenetic range (including batoids, galeomorphs, squalomorphs, and holocephalans; Fig. 1), as well as diverse body morphologies, habitats, and feeding modes. The ontogenetic stage of each individual was determined with aid of the IUCN Red List reports which detail the length at birth and maturity for each taxon. Because chondrichthyan diets frequently vary by location and age (Ebert et al., 2021; Last et al., 2016), each species was coded based on its sampling provenance and ontogenetic stage. For example, our *Scyliorhinus* specimen was collected in Algeria, so its assigned diet reflects dietary studies from that area of the Mediterranean. Ontogenetic dietary variation was likewise incorporated when known. Detailed prey items are provided in Table 1, and diets are categorised in the results as durophagous (predominantly hard-shelled prey), piscivorous (predominantly fish), or generalist/opportunistic (a mixed diet). Complete specimen details and references are available in Tables 1 and 2.

**Table 1.** General ecology information on the chondrichthyan taxa examined in this study. 1 (Lasso et al., 1996; Shibuya et al., 2009; Torres et al., 2025), 2: (Capapé et al., 2004; Kyne and Jabado, 2019), 3: (El Kamel-Moutalibi et al., 2013; Jabado et al., 2021a; Patokina and Litvinov, 2005), 4: (Jabado et al., 2021b; Patokina and Litvinov, 2005), 5: (Carlson et al., 2021; Motta et al., 2008), 6: (Jabado et al., 2021c; Smale and Compagno, 1997), 7: (Ba et al., 2013; Rigby et al., 2020; Sen et al., 2018), 8: (Finucci et al., 2021; Mnasri et al., 2012) 9:(Gordon et al., 2019; Morey et al., 2019), 10: (Finucci, 2020; Moura et al., 2005).

|  | **Scientific name** | **Specimen nos.** | **Diet** | **Depth range (m)** |
|---|---|---|---|---|
| 1 | *Glaucostegus cemiculus* | RMNH.PISC.27850 | Invertebrates, fish, cephalopods | 9–100 |
| 2 | *Potamotrygon orbignyi* | RMNH.PISC 37488 | Insect larvae, molluscs, crustaceans and fish | Freshwater rivers |
| 3 | *Torpedo torpedo* | RMNH.PISC.29360 | Fish, cephalopods | 2–400 |
| 4 | *Zanobatus schoenleinii* | ZMA.113.051 | Predominantly crustaceans, some cephalopods, fish and invertebrates | 10–15 (max ~100) |
| 5 | *Ginglymostoma cirratum* | ZMA 119.897 | Invertebrates, fish | 0–130 |
| 6 | *Mustelus Mustelus* | RMNH.PISC.34043 | Early ontogeny: crustaceans and invertebrates<br>Later ontogeny:<br>Cephalopods | 5–624 |
| 7 | *Rhizoprionodon acutus* | RMNH.PISC.38252 | Early ontogeny: Fish<br>Later ontogeny:<br>Fish, crustaceans and invertebrates | 1–200 |
| 8 | *Scyliorhinus canicula* | RMNH.PISC 15599 | Early ontogeny: fish crustaceans, cephalopods and invertebrates<br>Later ontogeny: Increase in crustaceans | 10–780 |
| 9 | *Squatina oculata* | ZMA 108.512 | Fish, cephalopods and crustaceans | 5–500 |
| 10 | *Chimaera monstrosa* | ZMA 112.915 | Early ontogeny: Invertebrates<br>Later ontogeny:<br>Crustaceans and invertebrates | 40–1400 |

**Table 2.**
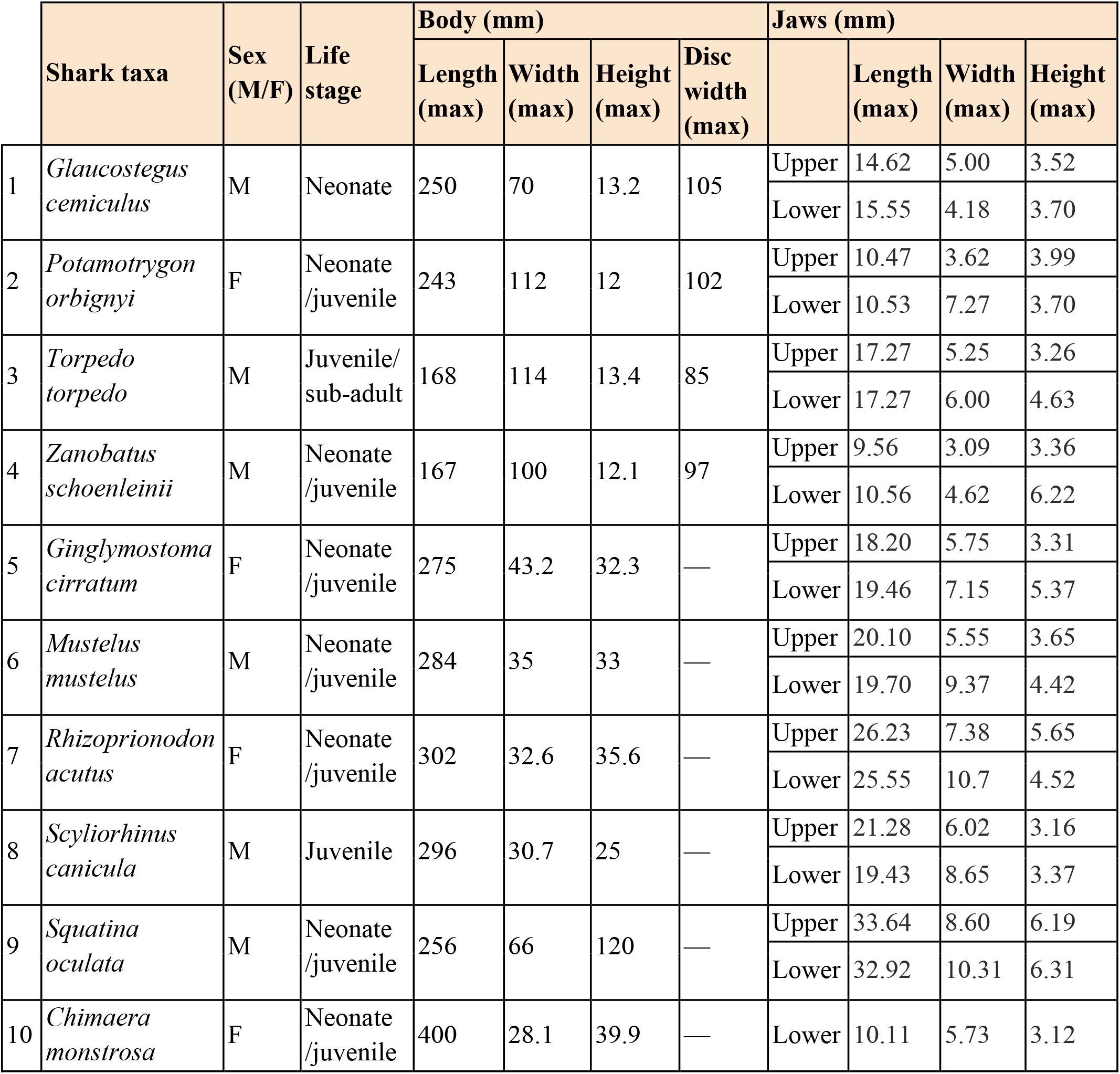
Specimen number and size data on chondrichthyan taxa used in this study.

|  | **Shark taxa** | **Sex (M/F)** | **Life stage** | **Body (mm)** |  |  |  | **Jaws (mm)** |  |  |  |
|---|---|---|---|---|---|---|---|---|---|---|---|
|  |  |  |  | **Length (max)** | **Width (max)** | **Height (max)** | **Disc width (max)** |  | **Length (max)** | **Width (max)** | **Height (max)** |
| 1 | *Glaucostegus cemiculus* | M | Neonate | 250 | 70 | 13.2 | 105 | Upper | 14.62 | 5.00 | 3.52 |
|  |  |  |  |  |  |  |  | Lower | 15.55 | 4.18 | 3.70 |
| 2 | *Potamotrygon orbignyi* | F | Neonate /juvenile | 243 | 112 | 12 | 102 | Upper | 10.47 | 3.62 | 3.99 |
|  |  |  |  |  |  |  |  | Lower | 10.53 | 7.27 | 3.70 |
| 3 | *Torpedo torpedo* | M | Juvenile/sub-adult | 168 | 114 | 13.4 | 85 | Upper | 17.27 | 5.25 | 3.26 |
|  |  |  |  |  |  |  |  | Lower | 17.27 | 6.00 | 4.63 |
| 4 | *Zanobatus schoenleinii* | M | Neonate /juvenile | 167 | 100 | 12.1 | 97 | Upper | 9.56 | 3.09 | 3.36 |
|  |  |  |  |  |  |  |  | Lower | 10.56 | 4.62 | 6.22 |
| 5 | *Ginglymostoma cirratum* | F | Neonate /juvenile | 275 | 43.2 | 32.3 | — | Upper | 18.20 | 5.75 | 3.31 |
|  |  |  |  |  |  |  |  | Lower | 19.46 | 7.15 | 5.37 |
| 6 | *Mustelus mustelus* | M | Neonate /juvenile | 284 | 35 | 33 | — | Upper | 20.10 | 5.55 | 3.65 |
|  |  |  |  |  |  |  |  | Lower | 19.70 | 9.37 | 4.42 |
| 7 | *Rhizoprionodon acutus* | F | Neonate /juvenile | 302 | 32.6 | 35.6 | — | Upper | 26.23 | 7.38 | 5.65 |
|  |  |  |  |  |  |  |  | Lower | 25.55 | 10.7 | 4.52 |
| 8 | *Scyliorhinus canicula* | M | Juvenile | 296 | 30.7 | 25 | — | Upper | 21.28 | 6.02 | 3.16 |
|  |  |  |  |  |  |  |  | Lower | 19.43 | 8.65 | 3.37 |
| 9 | *Squatina oculata* | M | Neonate /juvenile | 256 | 66 | 120 | — | Upper | 33.64 | 8.60 | 6.19 |
|  |  |  |  |  |  |  |  | Lower | 32.92 | 10.31 | 6.31 |
| 10 | *Chimaera monstrosa* | F | Neonate /juvenile | 400 | 28.1 | 39.9 | — | Lower | 10.11 | 5.73 | 3.12 |

### Micro-CT scanning and 3D modelling

Tesserae are typically hundreds of microns wide and thick (Maisey et al., 2020a; Seidel et al., 2020a), requiring high-resolution imaging techniques to visualise them (Baum et al., 2019; Knötel et al., 2017). Micro-CT scanning and 3D modelling workflows are well established for non-invasive, high-magnification 3D visualisations of mineralised tissues in both fossil and extant chondrichthyans (Clark et al., 2022; Jambura et al., 2019; Maisey et al., 2020a; Seidel et al., 2016; Stock et al., 2024). Specimens were scanned using the Zeiss X-radia 520 Versa scanner at Naturalis Biodiversity Center, the detailed scan parameters are listed in Table 3. The resolution of the reconstructed TIFF images (6 - 18.5 µm) allowed the TC to be clearly visualised in all but two taxa (*Mustelus mustelus*). The TC in *M.mustelus* was poorly resolved, which could be attributed to several factors, including scan resolution or prolonged specimen fixation. However, we ruled these possibilities out, as all other specimens were well-visualised despite prolonged fixation, with the TC clearly identifiable between 6 and 18.5 µm. We therefore concluded that the poor resolution was due to poorly developed TC in this specific specimen Reconstructed image stacks were imported into the 3D imaging software Materialise Mimics Core 26.0, where the upper and lower jaws were segmented based on grey-value thresholding using the mask function. Because contrast between the tesserae and the background was high, generating these masks was straightforward for most specimens. For *Squatina oculata*, in which mineralisation varied substantially within the tesseral layer, poorly mineralised areas were incorporated into the segmented volume using a second, lower-threshold mask. 3D models were generated from these masks and exported as STL files for rendering. Renders of the tesseral layer of the upper and lower jaws were generated using Blender 5.0. To visualise individual tesserae in detail, thick slabs were generated using VG Studio MAX 2024.4. For all taxa, multiple virtual thin sections were stacked to a thickness of 0.1 mm using a maximum intensity projection to highlight mineralised structures.

**Table 3.**
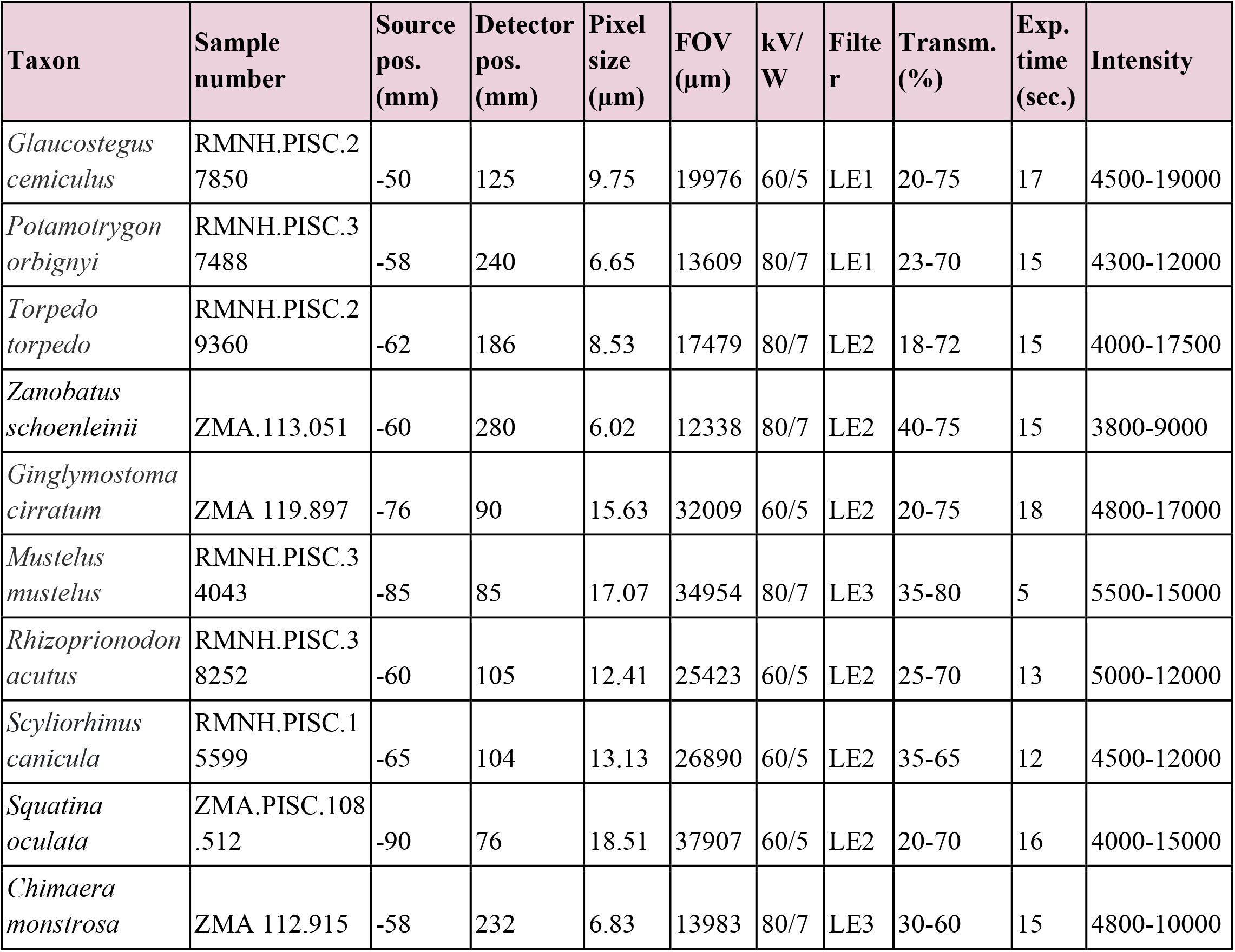
Scanning parameters for each specimen on a Zeiss Xradia 520 Versa 3D X-ray microscope.

| **Taxon** | **Sample number** | **Source pos. (mm)** | **Detector pos. (mm)** | **Pixel size (µm)** | **FOV (µm)** | **kV/W** | **Filter** | **Transm. (%)** | **Exp. time (sec.)** | **Intensity** |
|---|---|---|---|---|---|---|---|---|---|---|
| *Glaucostegus cemiculus* | RMNH.PISC.27850 | -50 | 125 | 9.75 | 19976 | 60/5 | LE1 | 20-75 | 17 | 4500-19000 |
| *Potamotrygon orbignyi* | RMNH.PISC.37488 | -58 | 240 | 6.65 | 13609 | 80/7 | LE1 | 23-70 | 15 | 4300-12000 |
| *Torpedo torpedo* | RMNH.PISC.29360 | -62 | 186 | 8.53 | 17479 | 80/7 | LE2 | 18-72 | 15 | 4000-17500 |
| *Zanobatus schoenleinii* | ZMA.113.051 | -60 | 280 | 6.02 | 12338 | 80/7 | LE2 | 40-75 | 15 | 3800-9000 |
| *Ginglymostoma cirratum* | ZMA 119.897 | -76 | 90 | 15.63 | 32009 | 60/5 | LE2 | 20-75 | 18 | 4800-17000 |
| *Mustelus mustelus* | RMNH.PISC.34043 | -85 | 85 | 17.07 | 34954 | 80/7 | LE3 | 35-80 | 5 | 5500-15000 |
| *Rhizoprionodon acutus* | RMNH.PISC.38252 | -60 | 105 | 12.41 | 25423 | 60/5 | LE2 | 25-70 | 13 | 5000-12000 |
| *Scyliorhinus canicula* | RMNH.PISC.15599 | -65 | 104 | 13.13 | 26890 | 60/5 | LE2 | 35-65 | 12 | 4500-12000 |
| *Squatina oculata* | ZMA.PISC.108.512 | -90 | 76 | 18.51 | 37907 | 60/5 | LE2 | 20-70 | 16 | 4000-15000 |
| *Chimaera monstrosa* | ZMA 112.915 | -58 | 232 | 6.83 | 13983 | 80/7 | LE3 | 30-60 | 15 | 4800-10000 |

### Tesserae measurements

Tesserae measurements were conducted using Materialise Mimics Core 26.0. For each scan, three small regions of interest (ROIs) were selected on both the lingual and labial surfaces at the proximal, middle, and distal portions of the jaws with a total of 12 regions per specimen (see Figure 2 for more details). These ROIs were positioned centrally between the oral and aboral margins and spaced evenly along the jaw’s length. Each ROI comprised ten clearly distinguishable tesserae, from which the axis with the **maximum width** (termed "**max width**" in the results) was recorded. Measurements across these six ROIs were then averaged to determine a mean tessera length per jaw. Specialised tesserae, such as the voussoir and columnar types, have not previously been studied during early ontogeny. Although we can assume that they will be more enlarged and more rectangular than standard tesserae, they are presumably not yet fully developed. To avoid misinterpretation, we measured the maximum width of tesserae which fit the criteria of being enlarged and rectangular, and provisionally termed them "specialised tesserae" in the results. Their specific morphological classifications are evaluated in the discussion. Tesseral layer thickness was quantified using the wall thickness function in Materialise 3-matic, which measures the distance between the external TC surface (perichondral) and internal TC surface (uncalcified cartilage) of a 3D mesh. Local thickness variation was visualised using false-colour heatmaps (e.g., Figure 4f, g), scaled in two ways: (1) a standardised range capped at 100 µm to allow direct comparisons across specimens (see SM figures for details), and (2) a specimen-specific range scaled to each specimen’s maximum thickness. Local variation in mineralisation is discussed not only in terms of tesseral layer thickness but also by noting regions with patchy tessellation or lower grey values in the scans (Figure 2). The absolute degree of mineralisation could not be quantified at a tissue-specific level (e.g., in mg HA/cm³), as the grey values in our datasets could not be standardised against known hydroxyapatite concentrations. As noted above, canal pores can be identified by their association with internal calcification (trabeculae). However, because fully formed trabeculae are not necessarily expected at early ontogenetic stages, their presence was not required to identify a canal pore. The maximum canal pore size was determined by measuring the widest axis using the measurement tools in Materialise Mimics Core 26.0.

**Figure 2.**
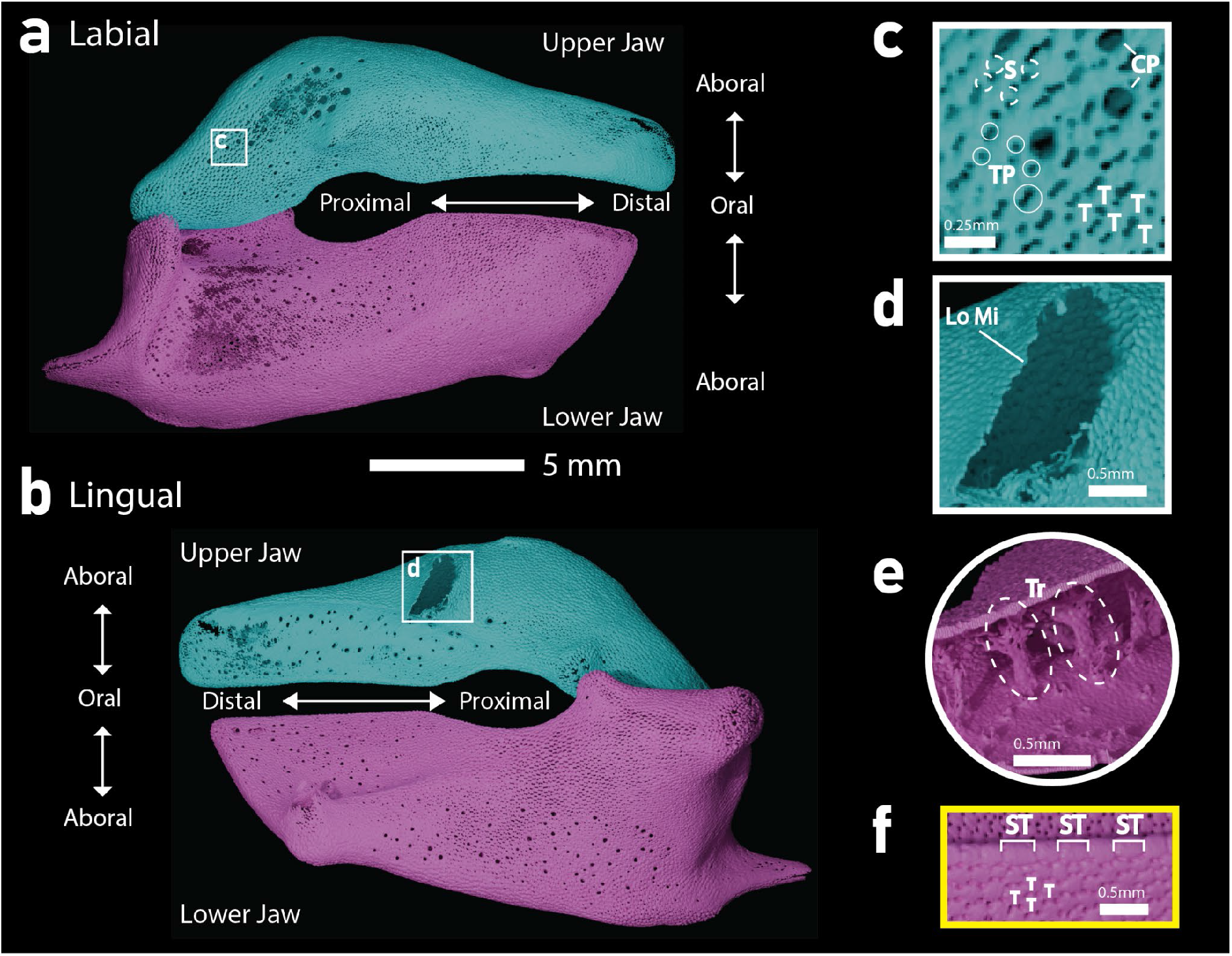
Directional terminologies and morphological features of Tessellated Cartilage (TC). (**a**) - (**e**) show the taxon *Ginglymostoma.* Labial (**a**) and lingual (**b**) views of the upper (turquoise) and lower (magenta) jaws with other directional terms labelled. Note that the teeth have been digitally segmented from the jaws and that the locations shown in **(c)**-**(d)** are indicated. The dashed circles in (**a**) and (**b**) show the approximate positions of the ROIs used for tesserae measurements. (**c**) Magnified region of the lower jaw with Tesserae (T), Spokes (S), Tesseral Pores (TP) and Canal Pores (CP) labelled. (**d**) Magnified region of no mineralisation from the upper jaw. (**e**) shows internal trabecular struts orientated lingual-labial at the proximal part of the lower jaw. (**f**) is highlighted in yellow as it is from the lower jaw of *Rhizoprionodon*, showing Tesserae (T) and Specialised Tesserae (ST).

## Results

### Batoids

#### Glaucostegus cemiculus

The tesserae are generally well-developed but their geometries (i.e. number of sides/neighbours) vary (Figure 3a). The average tessera max width for the upper jaw is 141 µm and the lower jaw 137 µm, giving an overall average of 139 µm (Table 3; Figure 3a and Figure 4a, d). The tesseral pores are notably large and the spokes are thin relative to the other batoids (Figure 3). There are small areas with low mineralisation at the articular surfaces labially (Figure 4b, e). The average tessera thickness is 40 µm, with a maximum thickness of 150 µm at the jaw socket of the lower jaw (Table 3; Figure 4f, g). There are numerous canal pores present on both jaws (the largest being 195 µm) which are difficult to differentiate from the large tesseral pores; they are visible in magnified zones (Figure 4e). These pores are associated with small internal mineralisations, signifying rudimentary trabeculae and thus we refer to them as canal pores (Table 4; Figure 2a, d, e). There is one complete trabecula that propagates across at the jaw joint socket of the lower jaw, with a length of 760 µm (Table 4; Figure 4c). There are no specialised tesserae present.

**Figure 3.**
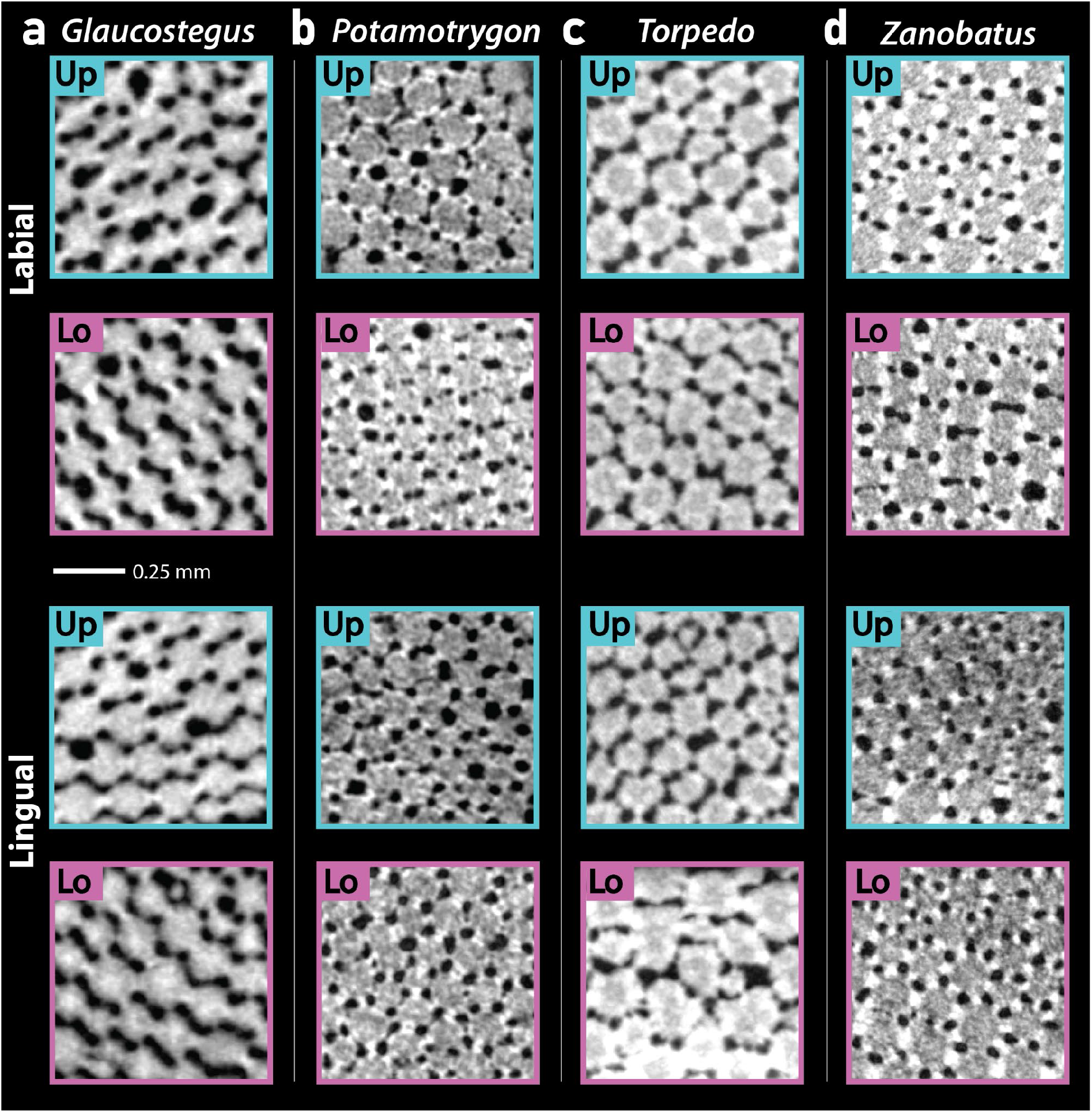
Virtual thick slabs of tesserae from the Upper (Up) and Lower (Lo) jaws of the four studied batoid taxa. **(a)** *Glaucostegus*, **(b)** *Potamotrygon*, **(c)** *Torpedo*, and **(d)** *Zanobatus* in labial and lingual view.

**Figure 4.**
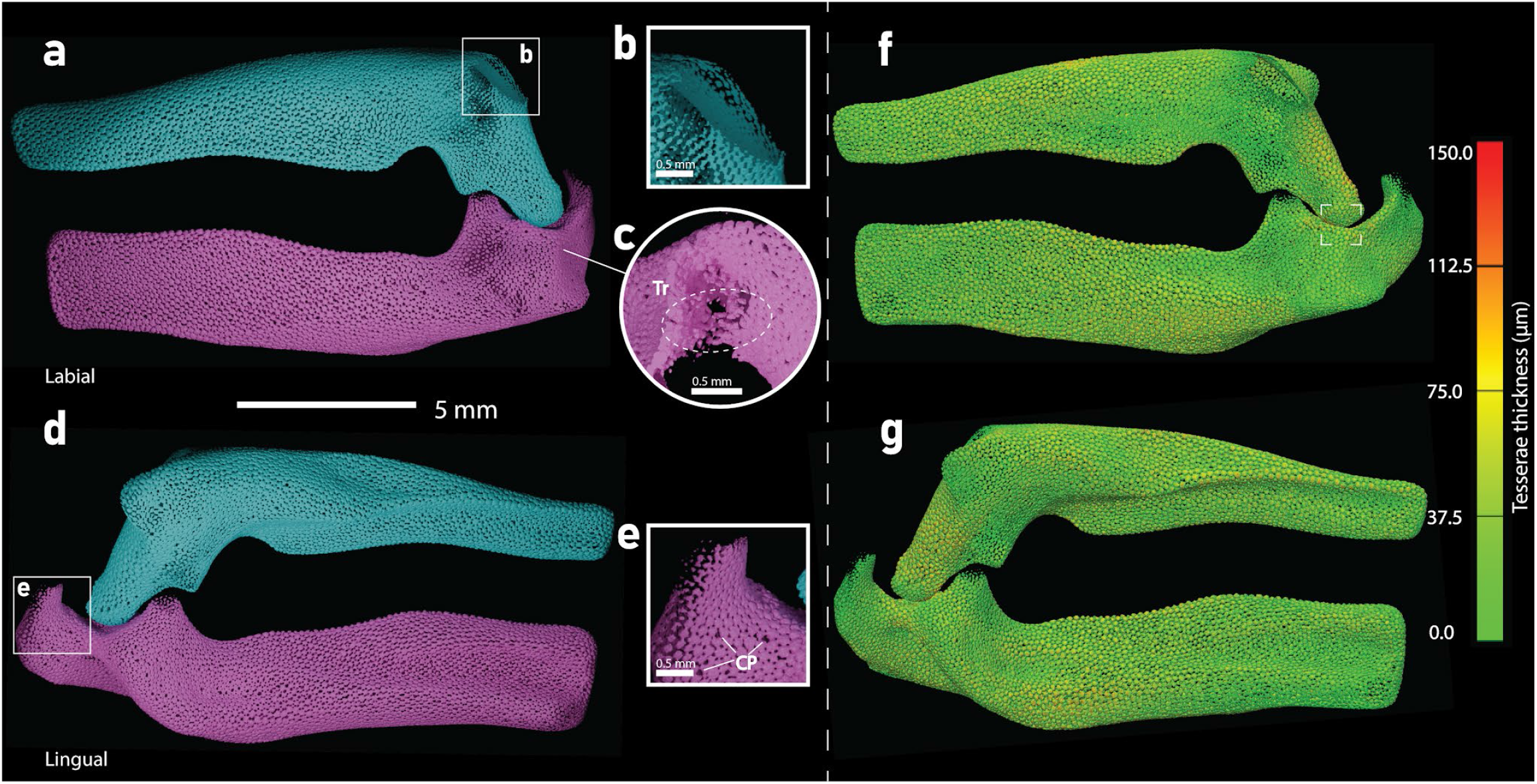
3D render of *Glaucostegus cemiculus* jaws. The upper jaw (turquoise) and lower jaw (magenta) are shown in labial **(a)** and lingual **(d)** views. Magnified regions of interest **(b, c, e)** highlighting Canal Pores (CP), trabeculae (Tr) and zones of lower mineralisation. Spatial thickness map of the TC is shown in labial (**f**) and lingual (**g**) view with the colour scale saturating at maximum thickness. The white box on panel (**f**) is where the thickest region is located

**Table 4.** Quantitative data on tessera size and tesseral layer thickness measurements for the upper and lower jaw for all specimens in this study.

| Scientific Name | Jaw | Labial surface average tessera length (µm) |  |  |  | Lingual surface average tessera length (µm) |  |  |  | Tessera length average (per jaw) | Overall tessera length average (µm) | Specialised Structures (µm) |  |  | Tesseral layer Thickness (µm) |  | Tessera max width/tessera thickness | Tessera max width/jaw length | Tessera max width/total body length |
|  |  | Anterior | Middle | Posterior | Lat Avg | Anterior | Middle | Posterior | Ling Avg |  |  | Specialised tesserae | Specialised tesserae | Trabeculae | Avg | Max |  |  |  |
|---|---|---|---|---|---|---|---|---|---|---|---|---|---|---|---|---|---|---|---|
| *Glaucostegus cemiculus* | Upper | 160± 14 | 153± 25 | 131± 22 | **148± 24** | 130± 32 | 155±20 | 130± 32 | **139±30** | **143±27** | **140±24** | *N/A* | *N/A* | *N/A* | 40 | 150 | 3.5 | 9.8 | 0.6 |
|  | Lower | 139± 16 | 151± 17 | 122± 24 | **138± 22** | 134± 23 | 145± 20 | 132± 19 | **137±21** | **138±21** |  | *N/A* | *N/A* | **760** |  |  |  | 8.9 |  |
| *Potamotrygon orbignyi* | Upper | 141± 19 | 121±24 | 157± 22 | **140±26** | 111±13 | 128± 14 | 143± 19 | **127±20** | **134±24** | **131± 26** | *N/A* | *N/A* | *N/A* | 17 | 70 | 7.7 | 12.8 | 0.5 |
|  | Lower | 129± 19 | 132± 28 | 128±17 | **130±21** | 106± 37 | 146± 36 | 135± 18 | **129±35** | **129±29** |  | *N/A* | *N/A* | **935** |  |  |  | 12.3 |  |
| *Torpedo torpedo* | Upper | 201± 26 | 158±21 | 199± 24 | **186±31** | 171± 25 | 128± 20 | 185± 24 | **161±33** | **174±34** | **174±35** | *N/A* | *N/A* | *N/A* | 39 | 140 | 4.5 | 10.1 | 1 |
|  | Lower | 190± 41 | 181± 36 | 171± 34 | **181±37** | 149± 26 | 148± 14 | 212± 17 | **170±36** | **175±36** |  | *N/A* | *N/A* | *N/A* |  |  |  | 10.1 |  |
| *Zanobatus schoenleinii* | Upper | 142± 11 | 147± 21 | 133± 23 | **141±19** | 132± 10 | 137± 23 | 171± 23 | **146±26** | **144±23** | **146±24** | 159 | **184** | *N/A* | 18 | 80 | 8.1 | 15.1 | 0.9 |
|  | Lower | 152± 22 | 149±35 | 151±20 | **151±26** | 127± 27 | 159±18 | 152± 15 | **146±24** | **148±25** |  | 209 |  | *N/A* |  |  |  | 14.0 |  |
| *Ginglymostoma cirratum* | Upper | 119± 18 | 115± 14 | 138± 22 | **119± 18** | 134± 27 | 140± 19 | 135± 8 | **137±19** | **130±20** | **131±22** | *N/A* | *N/A* | ***740*** | 49 | 180 | 2.7 | 7.1 | 0.5 |
|  | Lower | 148± 21 | 115± 20 | 124± 19 | **148± 21** | 124± 20 | 138±21 | 147± 16 | **129±24** | **132±23** |  | *N/A* | *N/A* | ***1025*** |  |  |  | 6.8 |  |
| *Mustelus mustelus* | Upper | *N/A* | *N/A* | *N/A* | N/A | *N/A* | *N/A* | *N/A* | N/A | N/A | N/A | *N/A* | *N/A* | *N/A* | 41 | 130 | N/A | *N/A* | N/A |
|  | Lower | *N/A* | *N/A* | *N/A* | N/A | *N/A* | *N/A* | *N/A* | N/A | N/A |  | *N/A* | *N/A* | *N/A* |  |  |  | *N/A* |  |
| *Rhizoprionodon acutus* | Upper | 168± 33 | 244± 42 | 229± 33 | **214±48** | 165±23 | 171±32 | 160± 38 | **165±31** | **189±47** | **184±44** | 242 | ***270*** | *N/A* | 55 | 250 | 3.3 | 7.2 | 0.6 |
|  | Lower | 220± 17 | 169± 39 | 152± 56 | **180±49** | 172± 36 | 190± 19 | 163± 29 | **175±30** | **178±40** |  | 297 |  | *N/A* |  |  |  | 7.0 |  |
| *Scyliorhinus canicula* | Upper | 145± 19 | 130± 18 | 184± 40 | **153±35** | 135±15 | 129± 28 | 174±25 | **146±30** | **150±33** | **145±31** | 187 | ***204*** | *N/A* | 33 | 200 | 4.4 | 7.0 | 0.5 |
|  | Lower | 158± 20 | 141± 17 | 177± 17 | **158±23** | 141± 16 | 114± 21 | 112± 13 | **122±21** | **140±29** |  | 220 |  | *N/A* |  |  |  | 7.2 |  |
| *Squatina oculata* | Upper | 272± 29 | 297± 26 | 262± 37 | **277±33** | 256±38 | 359± 61 | 270± 24 | **295±63** | **286±51** | **270±48** | *N/A* | *N/A* | *N/A* | 63 | 7.7 | 4.3 | 8.5 | 1.1 |
|  | Lower | 240± 37 | 272± 42 | 249± 49 | **254±44** | 248± 28 | 255± 39 | 254±34 | **252±33** | **253±38** |  | *N/A* |  | *N/A* |  |  |  | 7.7 |  |
| *Chimaera monstrosa* | Lower | *N/A* | *N/A* | *N/A* | N/A | *N/A* | *N/A* | *N/A* | N/A | N/A | N/A | *N/A* | *N/A* | *N/A* | 17 |  | *N/A* | *N/A* | *N/A* |

#### Potamotrygon orbignyi

The tesserae are well-developed, polygonal, and generally homogeneous in size and shape (Figure 3b). The average max tessera width differs by just 2 µm between the upper and lower jaws, with a mean max width of 132 µm across both jaws (Table 4; Figures 3b, 5a, b, d). Tesserae are relatively thin, with an average thickness of 17 µm and a maximum thickness of 130 µm along the oral margin of the upper jaw (Table 4; Figure 5g, h). Specialised tesserae with an average width of 156 µm line the narrow oral margins at the distal ends of both jaws (Figure 5a, b, d). Small regions of lower mineralisation occur at the symphysis and jaw articular surfaces (Figure 5a, d, f). Numerous canal pores are present on both lateral and lingual surfaces, with the largest reaching 161µm (Table 4; Figure 5a, d, e). Internally, these openings are associated with rudimentary trabeculae (hence our designation as "canal pores"), with a single complete trabecula measuring 935 µm in length located posteriorly on the lower jaw (Table 4; Figure 5c). *Torpedo torpedo*

**Figure 5.**
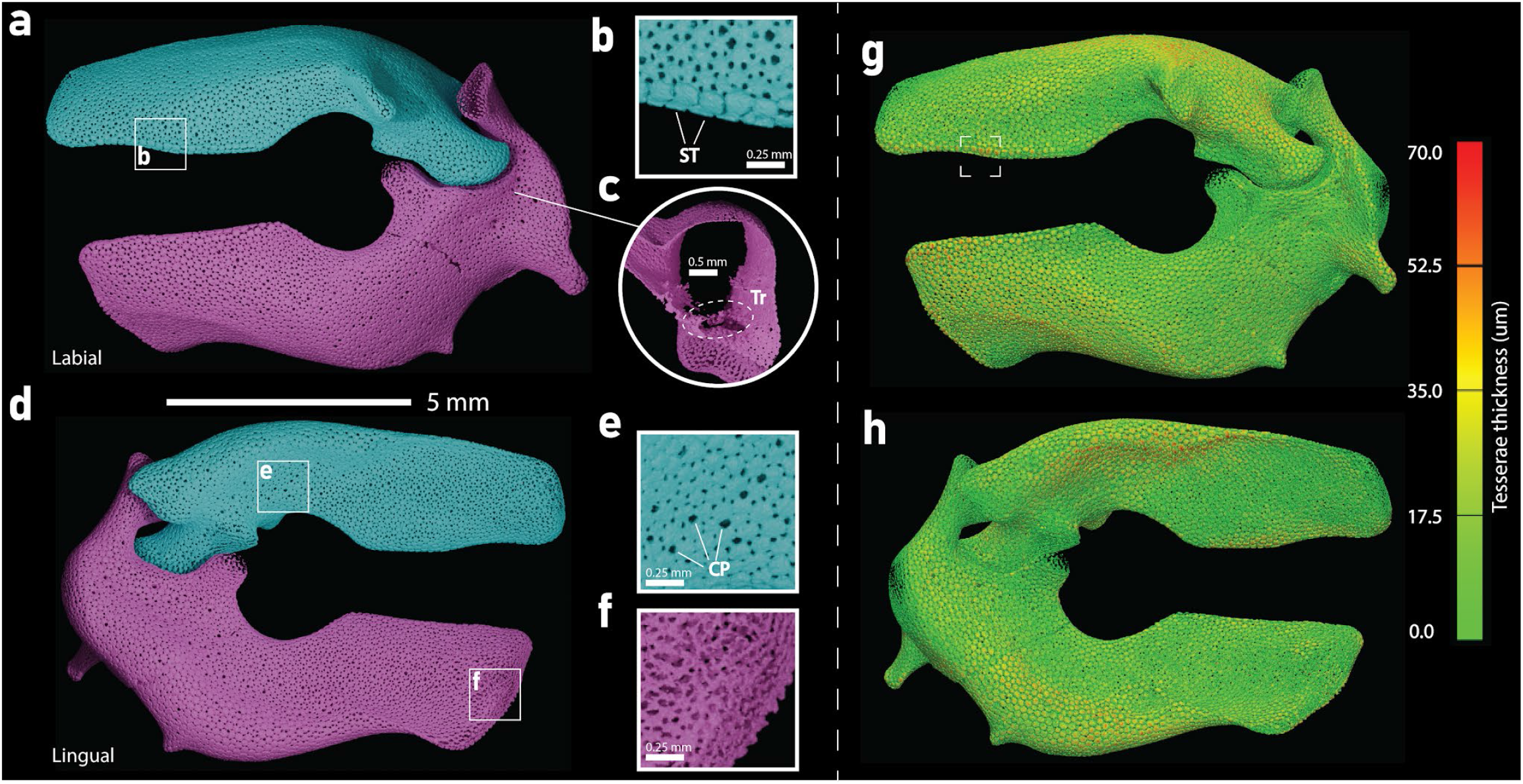
3D render of *Potamotrygon orbignyi* jaws. The upper jaw (turquoise) and lower jaw (magenta) are shown in labial **(a)** and lingual **(d)** views. Magnified regions of interest **(b, e, f)** highlighting Canal Pores (CP), Specialised Tesserae (ST) and zones of lower mineralisation. Section of lower jaw (**c**) showing trabeculae (Tr). Spatial thickness map of the TC is shown in labial (**g**) and lingual (**h**) view with the colour scale saturating at maximum thickness. The white box on panel (**g**) is where the thickest region is located

The tesserae are generally polygonal with short and thin spokes in between, relative to the other batoids in this study, seen clearly in the thick slabs (Figures 3). The tesserae have the same average max width for both jaws of 174 µm (Table 4). Tesserae thickness appears to be higher on the labial surface from the spatial thickness map, compared with the lingual, with an average tesserae thickness is 39 µm across both jaws with a maximum thickness of 140 µm along the aboral margin of the lower jaw (Table 4; Figure 6g, h). Small areas of lower mineralisation are found at the jaw articular surfaces (Figure 6b, e). There are no specialised tesserae, no canal pores nor any internal mineralised structures.

**Figure 6.**
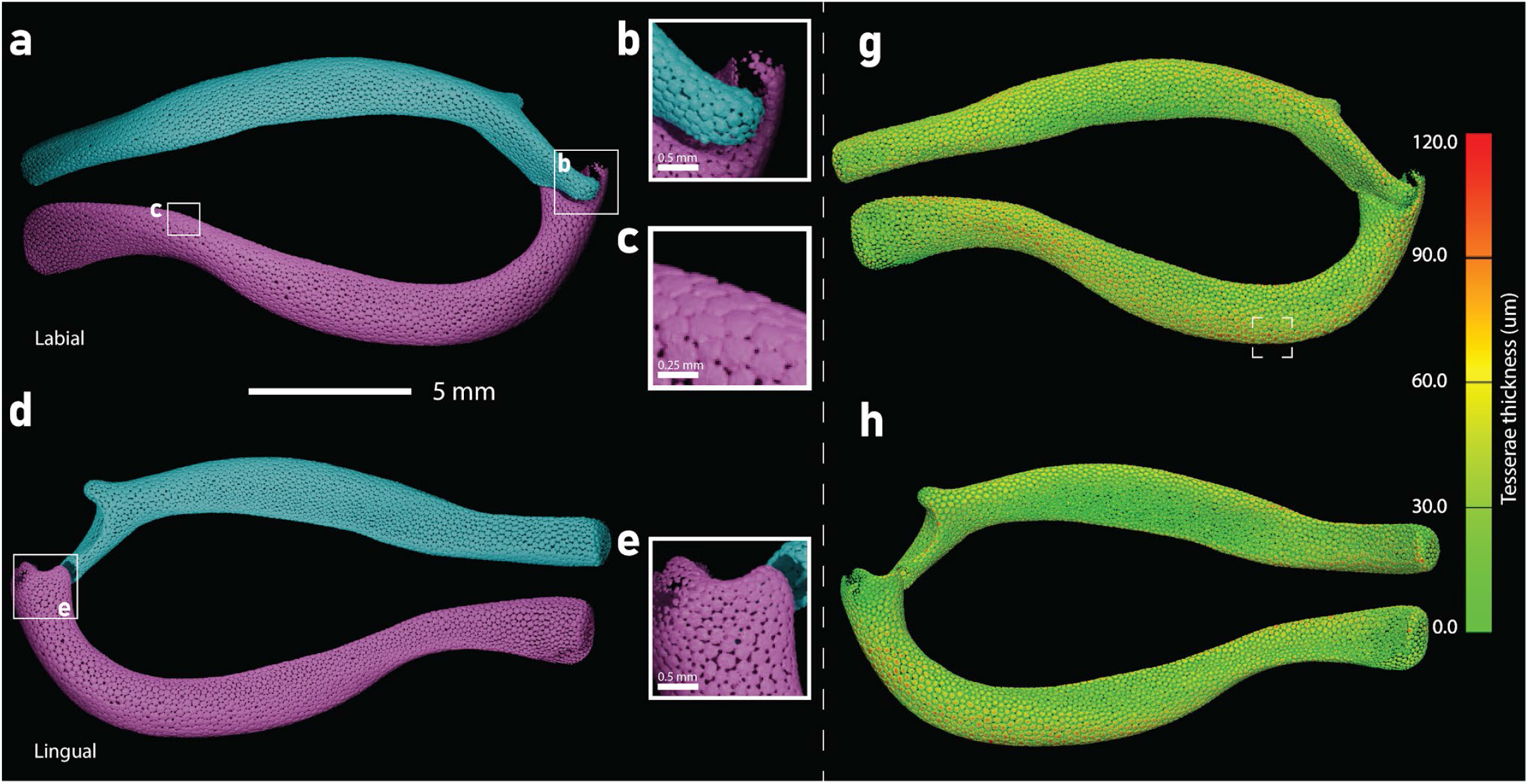
3D render of *Torpedo torpedo* jaws. The upper jaw (turquoise) and lower jaw (magenta) are shown in labial **(a)** and lingual **(d)** views. Magnified regions of interest **(b, c, e,)** highlighting zones of higher and lower mineralisation. Spatial thickness map of the TC is shown in labial (**g**) and lingual (**h**) view with the colour scale saturating at maximum thickness. The white box on panel (**g**) is where the thickest region is located

#### Zanobatus schoenleinii

The tesserae are well-developed and are polygonal, with small intertesseral spacing and thicker spokes compared with the other batoids seen which is best in the thick slabs (Figure 3 and Figure 7a, d). The average tessera max width for the upper jaw is 144 µm and 147 µm for the lower jaw, giving an overall average of 146 µm (Table 4). *Zanobatus* has the largest tesserae relative to jaw length of all the taxa in this study (Table 4) The average tesserae thickness is 18 µm, with a maximum thickness of 80 µm found on the lingual surface of the upper jaw (Table 4; Figure 7g, h). There are a couple of areas with lower mineralisation at the symphysis and jaw joint articulation (Figure 7b, c, f). Numerous large pores are present on both jaws, the majority located at the proximal end of the jaws (Figure 7a, d, e). The largest is 188 µm, larger than the average tesserae size (Table 4). These pores are associated with small protrusions of mineralisation, likely rudimentary trabeculae, and thus are referred to as canal pores. No specialised tesserae are observed.

**Figure 7.**
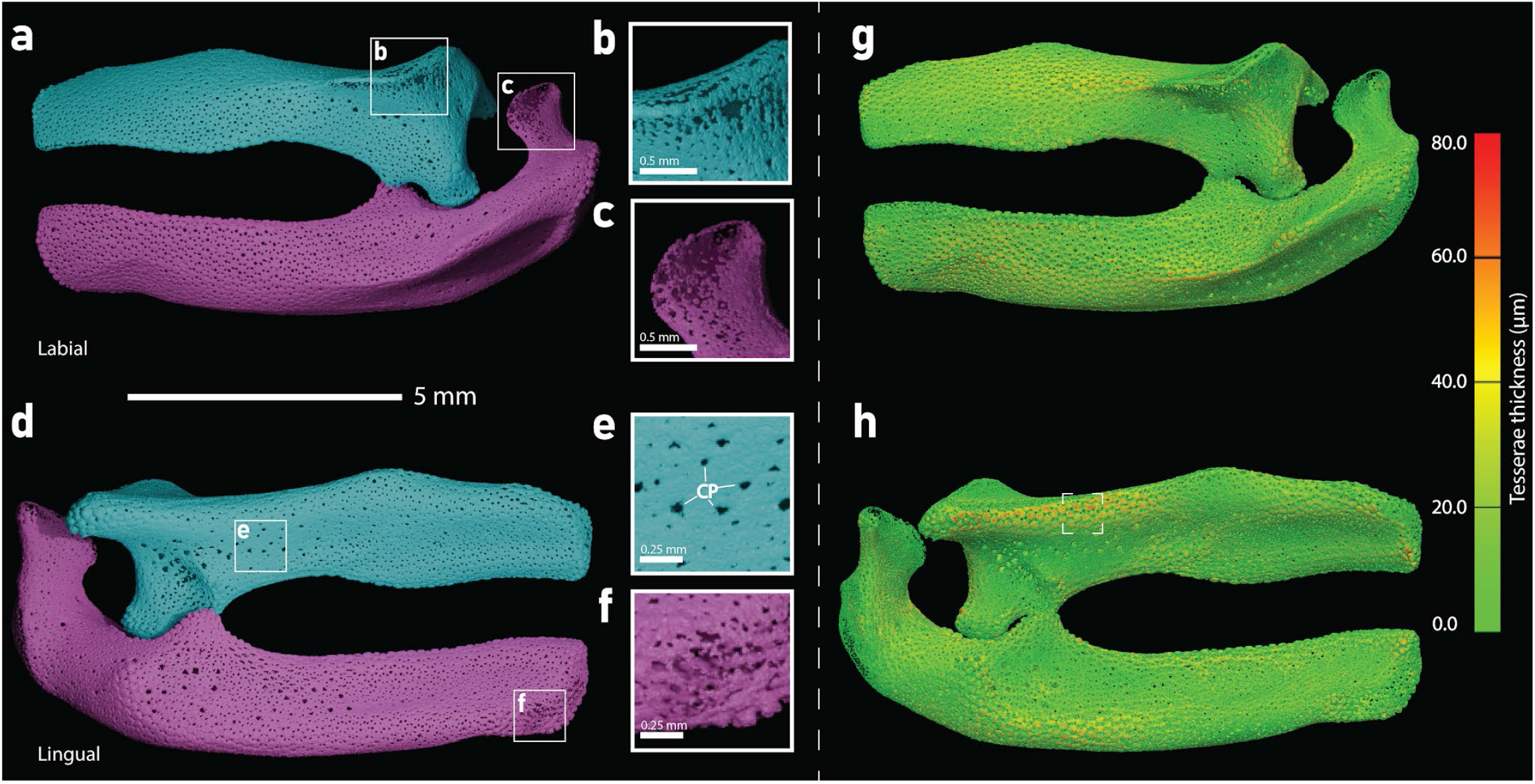
3D render of *Zanobatus schoenleinii* jaws. The upper jaw (turquoise) and lower jaw (magenta) are shown in labial **(a)** and lingual **(d)** views. Magnified regions of interest **(b, c, e, f)** highlighting Canal Pores (CP) and zones of lower mineralisation. Spatial thickness map of the TC is shown in labial (**g**) and lingual (**h**) view with the colour scale saturating at maximum thickness. The white box on panel (**h**) is where the thickest region is located

### Galeomorphs

#### Ginglymostoma cirratum

There is a high degree of heterogeneity in tesserae shape, size and mineralisation across both the upper and lower jaws, a departure from observations of the tesseral layer in the batoids (Figures 3, 8a and 9 a,d). Tesserae are most discernible in the central regions of the jaws, and they vary in shape with large tesseral pores and narrow spokes as seen in the thick slabs (Table 4; Figure 8a). Along the aboral margins, tesseral pores are absent and thus appear to fuse together (Figures 8a and 9a, f). The average tessera max. width for the upper jaw is 130 µm and 133 µm for the lower jaw, giving an overall average size of 132 µm, which is among the smallest tesserae of the taxa in this study (Table 4; SM Figures 11-13). The proximal ends of the jaws exhibit large areas of low mineralisation along their lateral surfaces, as well as on the lingual surface of the upper jaw. Mineralisation is low at the orbital process on the lingual surface of the upper jaw (Figure 9f). The TC thickness varies greatly across the jaws, with several regions exceeding 90 µm mostly in the aboral regions (Figure 9g,h; SM figure 5). The average thickness is 49 µm and a maximum thickness of 180 µm found on the aboral margin of the upper jaw (Table 4). Numerous canal pores are present (associated with trabeculae) with the largest being 520 µm, almost 4 times the average tessera size (Table 4; Figure 9b, c). On the upper jaw, canal pores can be found on the lingual surface near the teeth and proximally on the lateral surface (Figure 9a-f). On the lower jaw, they appear throughout the lingual surface (Figure 9e) and are concentrated towards the posterior end of the lateral surface, with the largest canal pores occurring near the jaw joint (Figure 9c).

**Figure 8.**
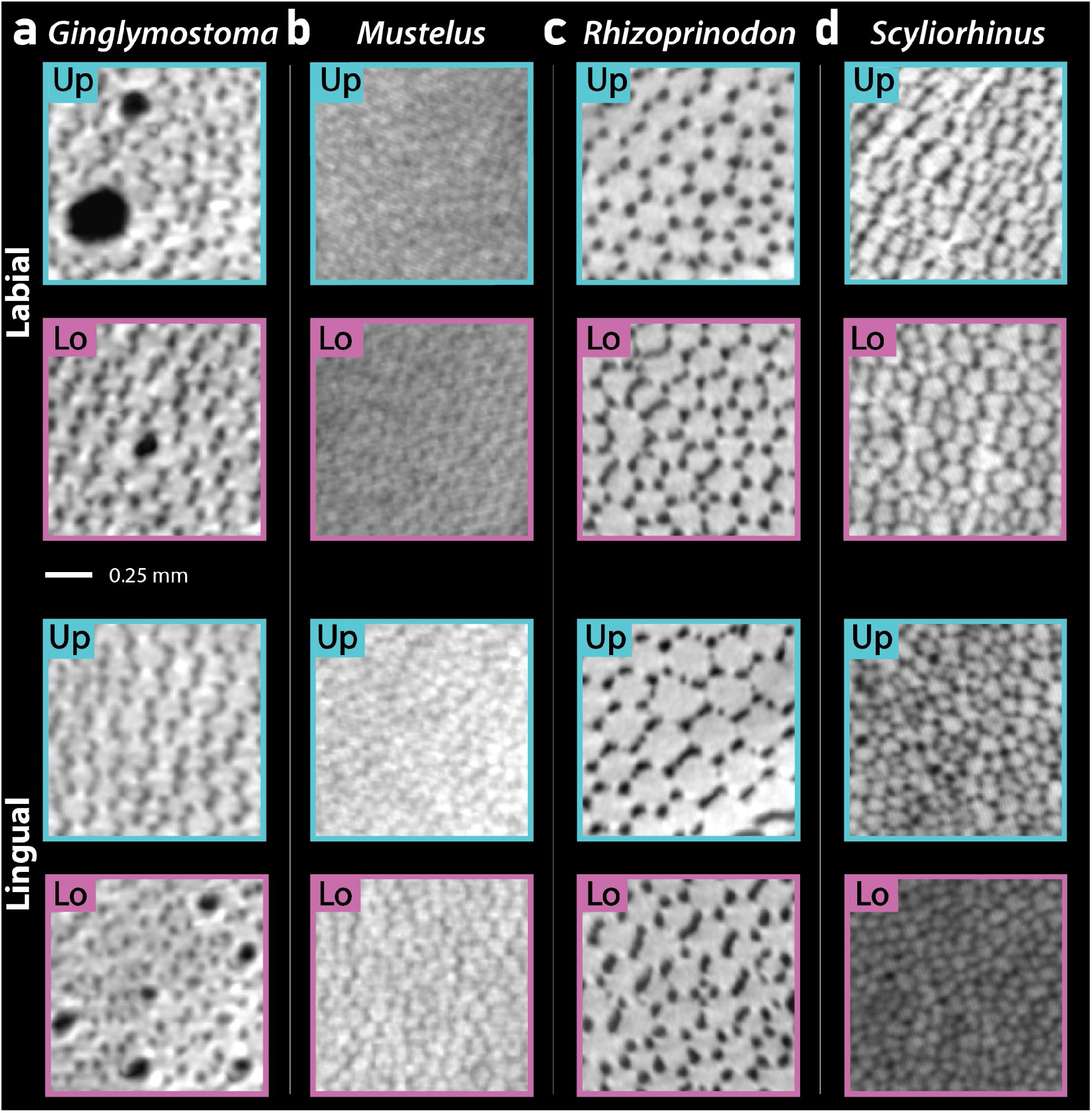
Virtual thick slabs of tesserae from the Upper (Up) and Lower (Lo) jaws of the five studied galeomorph taxa. **(a)** *Ginglymostoma*, **(b)** *Mustelus,* **(c)** *Rhizoprionodon* and **(d)** *Scyliorhinus* in labial and lingual view.

**Figure 9.**
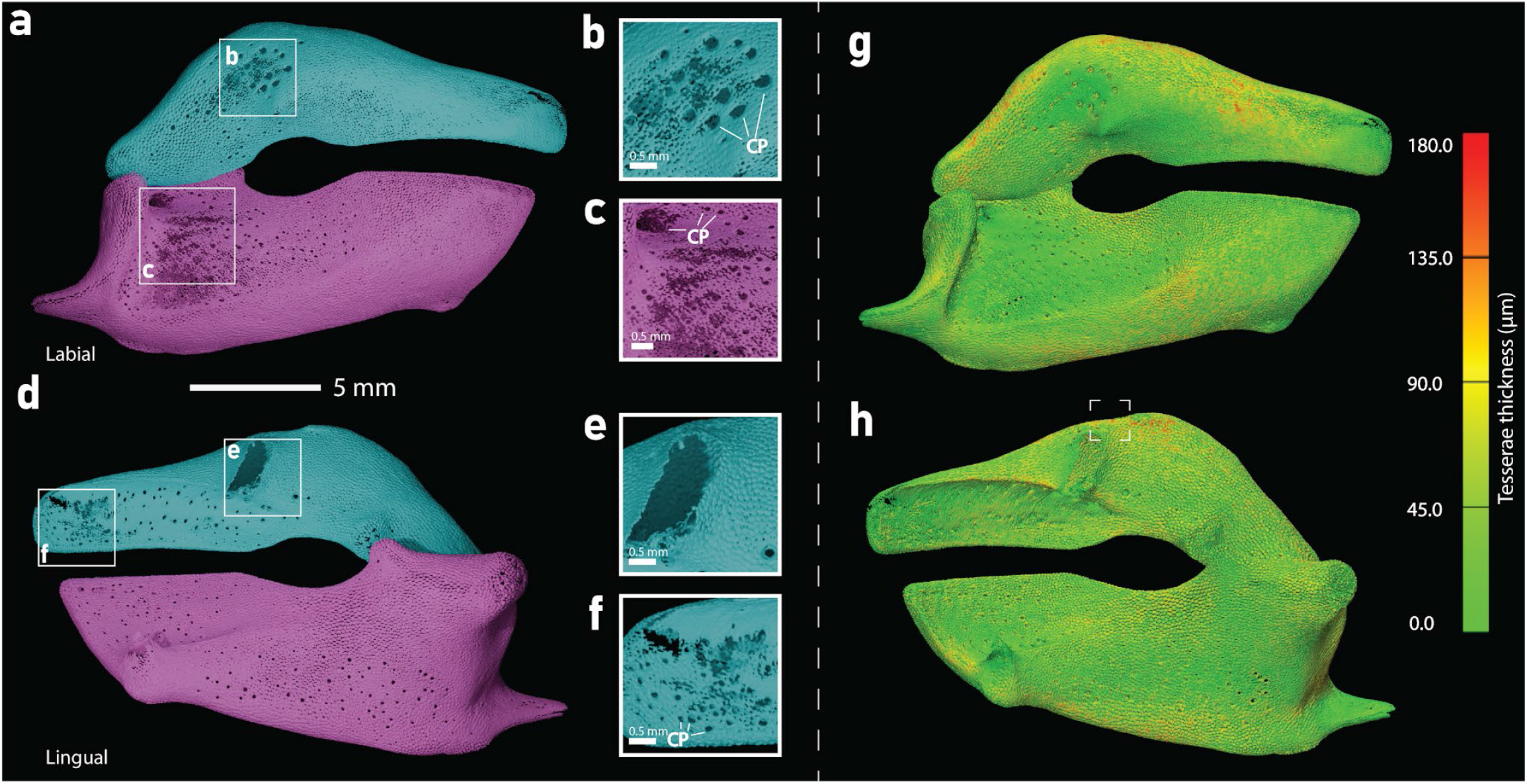
3D render of *Ginglymostoma cirratum* jaws. The upper jaw (turquoise) and lower jaw (magenta) are shown in labial **(a)** and lingual **(d)** views. Magnified regions of interest **(b, c, e, f)** highlighting Canal Pores (CP) and zones of lower mineralisation. Spatial thickness map of the TC is shown in labial (**g**) and lingual (**h**) view with the colour scale saturating at maximum thickness. The white box on panel (**h**) is where the thickest region is located.

Internally, trabeculae are numerous inside both jaws (Figure 10a-d). They vary in their degrees of development, but there are several that propagate fully between lingual and labial tesseral layers. Trabeculae in the upper jaw are mostly concentrated in the distal end, with the opposite being true for the lower jaw (Figure 10a-d). The majority of the trabeculae bridge labial-lingual sides, with a large network of trabeculae in the proximal end of the lower jaw propagating in multiple directions (Figure 10c).

**Figure 10.**
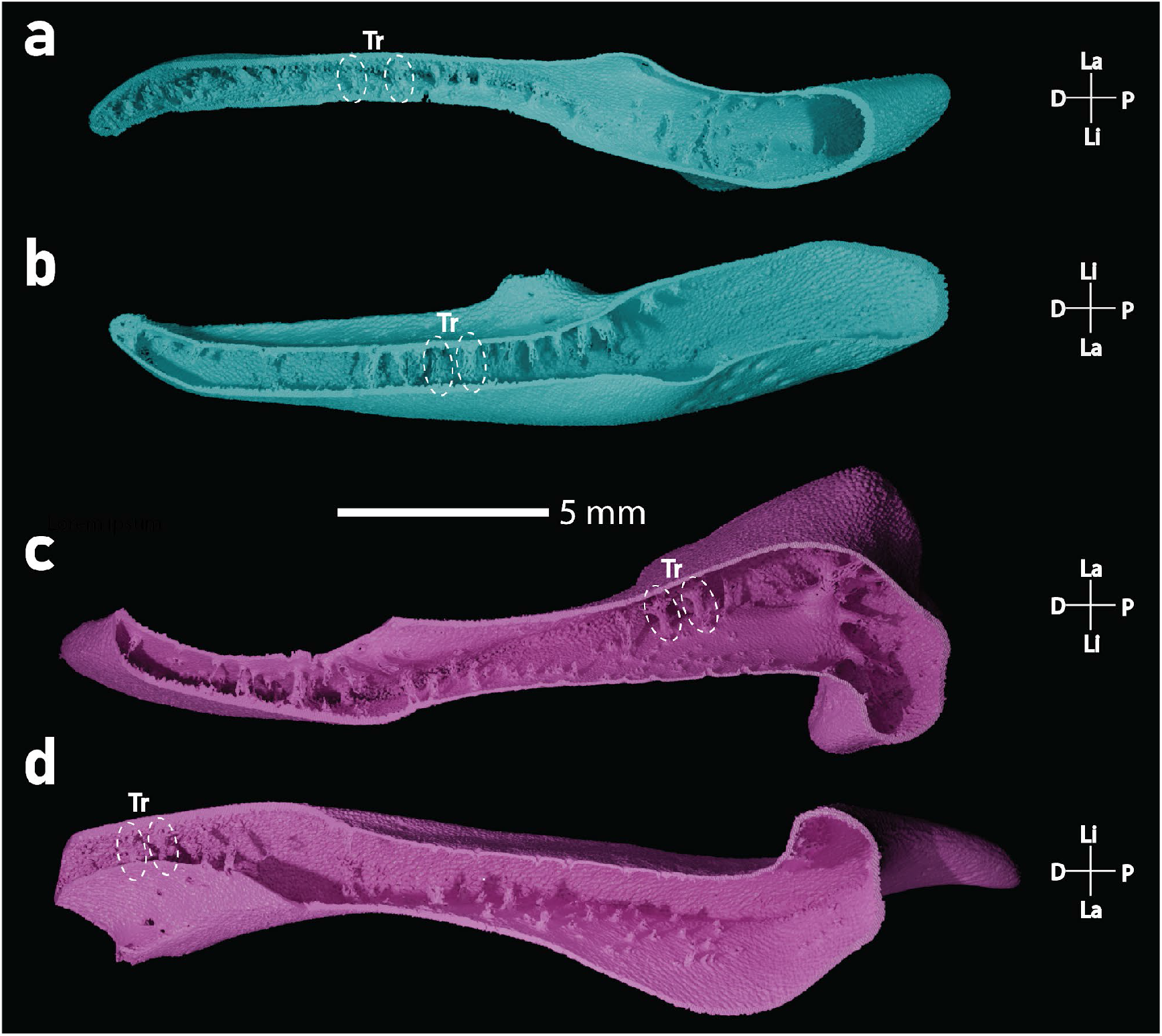
3D renders of the internal jaw morphology of *G. cirratum*. **(a, b)** Upper jaw (turquoise) and **(c, d)** lower jaw (magenta), showing internal trabeculae struts (Tr). Views include: **(a)** dorsal half, ventral view; **(b)** ventral half, dorsal view; **(c)** dorsal half, ventral view; and **(d)** ventral half, dorsal view. Axis key: D, distal; La, labial; Li, lingual; P, proximal.

#### Mustelus mustelus

Individual tesserae are faintly visible in the thick slabs, particularly on the lower jaw, whereas in the 3D models the tesseral layer appears as a homogeneous sheet of mineralisation (Figures 8b, 11a, d). The tesserae appear granular and lack discernible tesseral pores and spokes; due to this poor development, individual tessera dimensions were not measured (Figure 8b). A few large gaps occur in the tesseral layer, mostly on the lower jaw and around the jaw joints (Figure 11a–f). Despite the poorly developed tessellated cartilage, the tesseral layer is relatively thick compared to that of other taxa in this study (Table 4). Its thickness exceeds 65 µm across large regions of the labial surface (Figures 11g, h; Supplementary Figure 6) and averages 41 µm, reaching a maximum thickness of 130 µm on the labial surface near the distal end of the lower jaw (Table 4). Specialised tesserae, canal pores and internal mineralised structures are absent.

**Figure 11.**
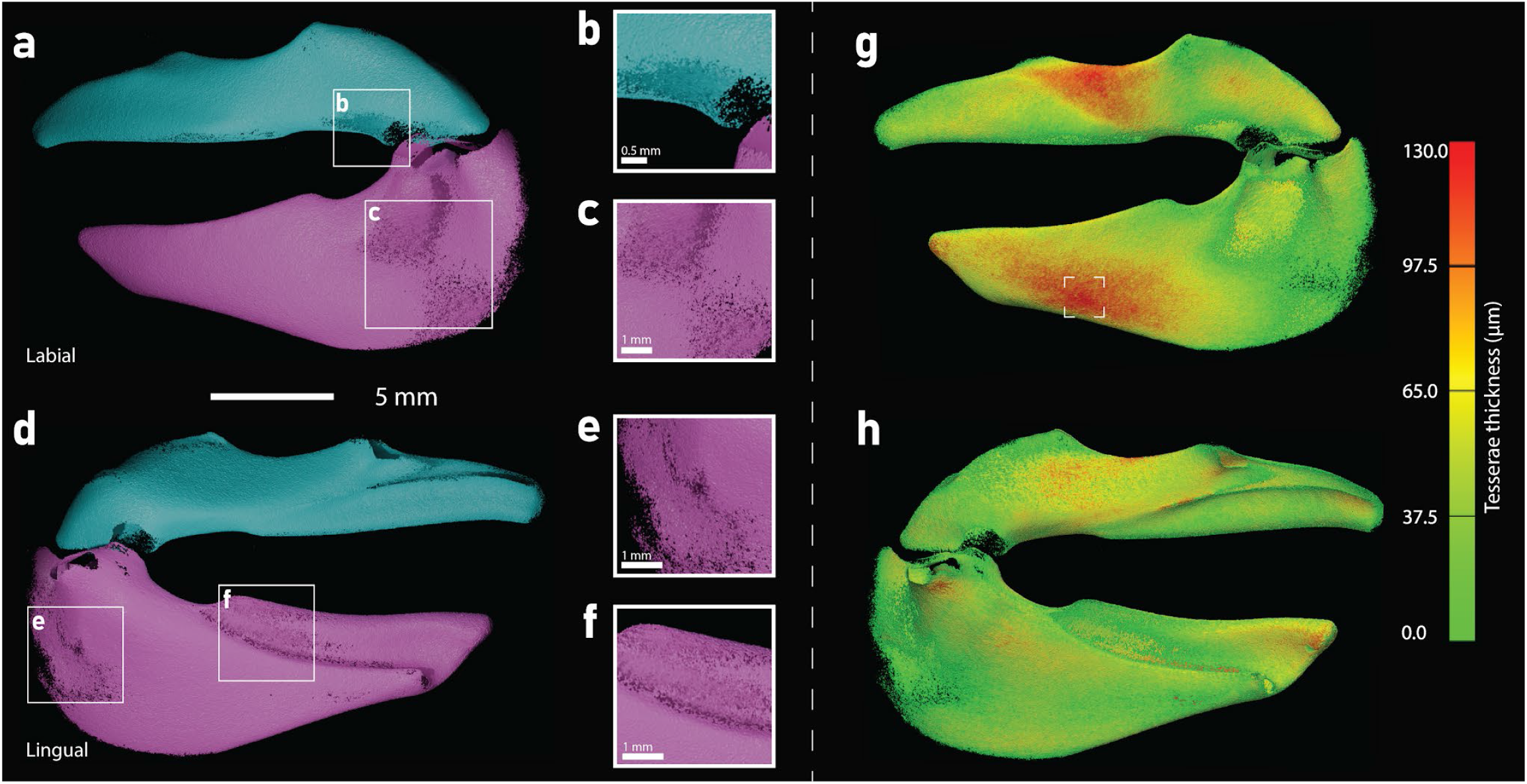
3D render of *Mustelus mustelus* jaws. The upper jaw (turquoise) and lower jaw (magenta) are shown in labial **(a)** and lingual **(d)** views. Magnified regions of interest **(b, c, e, f)** highlight zones of lower mineralisation. Spatial thickness map of the TC is shown in labial (**g**) and lingual (**h**) view with the colour scale saturating at maximum thickness. The white box on panel (**g**) is where the thickest region is located

#### Rhizoprionodon acutus

The tesserae are well-developed and homogeneous in shape and size with robust intertesseral spokes, which is reminiscent of the tessellated cartilage in the batoids (Figures 3, 8c and 12a, d). The average max tessera width is 189 µm on the upper jaw and 179 µm on the lower jaw, resulting in an overall average of 184 µm (Table 4; Figure 8c and Figure 12a, d). The tesseral layer is mostly complete, with small patches of low mineralisation at the orbital process on the upper jaw and proximally on the lower jaw on the labial surface (Figure 12c, e). The thickness of the tesseral layer exceeds 125 µm in several regions, notably the specialised tesserae and the distal ends of the labial surface (Figure 12g,h; SM figure 7). The average thickness is 55 µm with a maximum of 250 µm in the specialised tesserae of the lower jaw (Table 4; Figure 12g,h). There are specialised tesserae present on the oral margins of the upper and lower jaw, and they are present on an additional parallel margin on the lingual side of both jaws and they have an average max. width of 270 µm (Figure 12b,c,f). There are no canal pores present, and no mineralised structures present internally.

**Figure 12.**
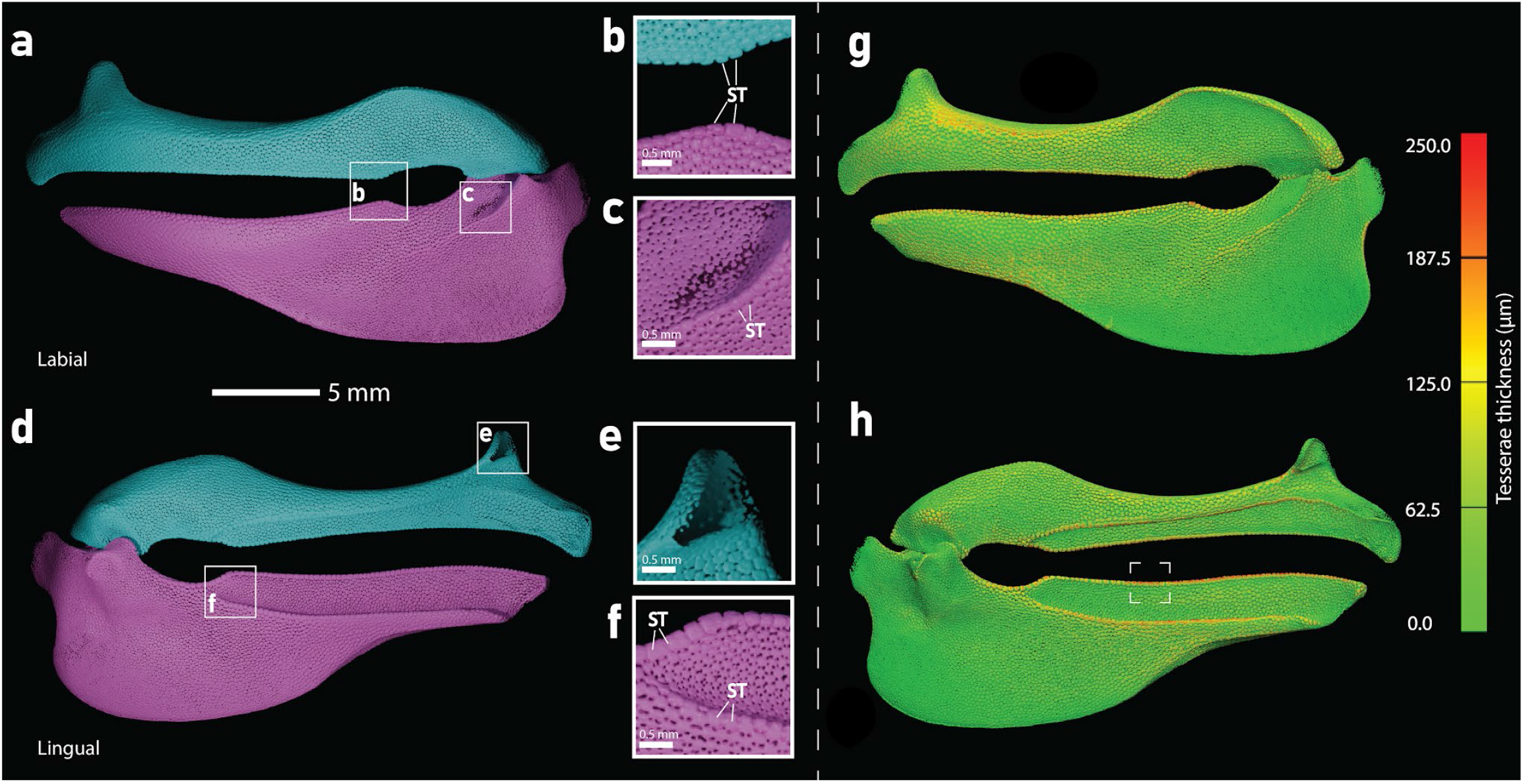
3D render of *Rhizoprionodon acutus* jaws. The upper jaw (turquoise) and lower jaw (magenta) are shown in labial (a) and lingual (d) views. Magnified regions of interest **(b, c, e, f)** highlight Specialised Tesserae (ST) and zones of lower mineralisation. Spatial thickness map of the TC is shown in labial (**g**) and lingual (**h**) view with the colour scale saturating at maximum thickness. The white box on panel (**h**) is where the thickest region is located.

#### Scyliorhinus canicula

There is a high degree of heterogeneity in tessera shape, size, and mineralisation across both the upper and lower jaws (Figures 8d, 13a, d). The tesserae are generally poorly developed, and are amorphous in shape; their appearance varies from fused with few intertesseral spaces to lacking spokes altogether (Figures 8d, 13a, d). Average tessera max width is 146 µm for the upper jaw and 142 µm for the lower jaw, yielding an overall mean of 144 µm (Table 4). Specialised tesserae are present along the oral margins beneath the teeth, with an average size of 204 µm. Several regions exhibit low or absent mineralisation, notably at the symphysis, jaw joints, ethmoid process of the upper jaw, and aboral region of the lower jaw (Figure 13a, b, d–f). The tesseral layer reaches its maximum thickness (200 µm) in the specialised tesserae of the lower jaw, with an overall average thickness of 33 µm (Table 4; Figure 13g, h). Canal pores and internal mineralised structures are absent.

**Figure 13.**
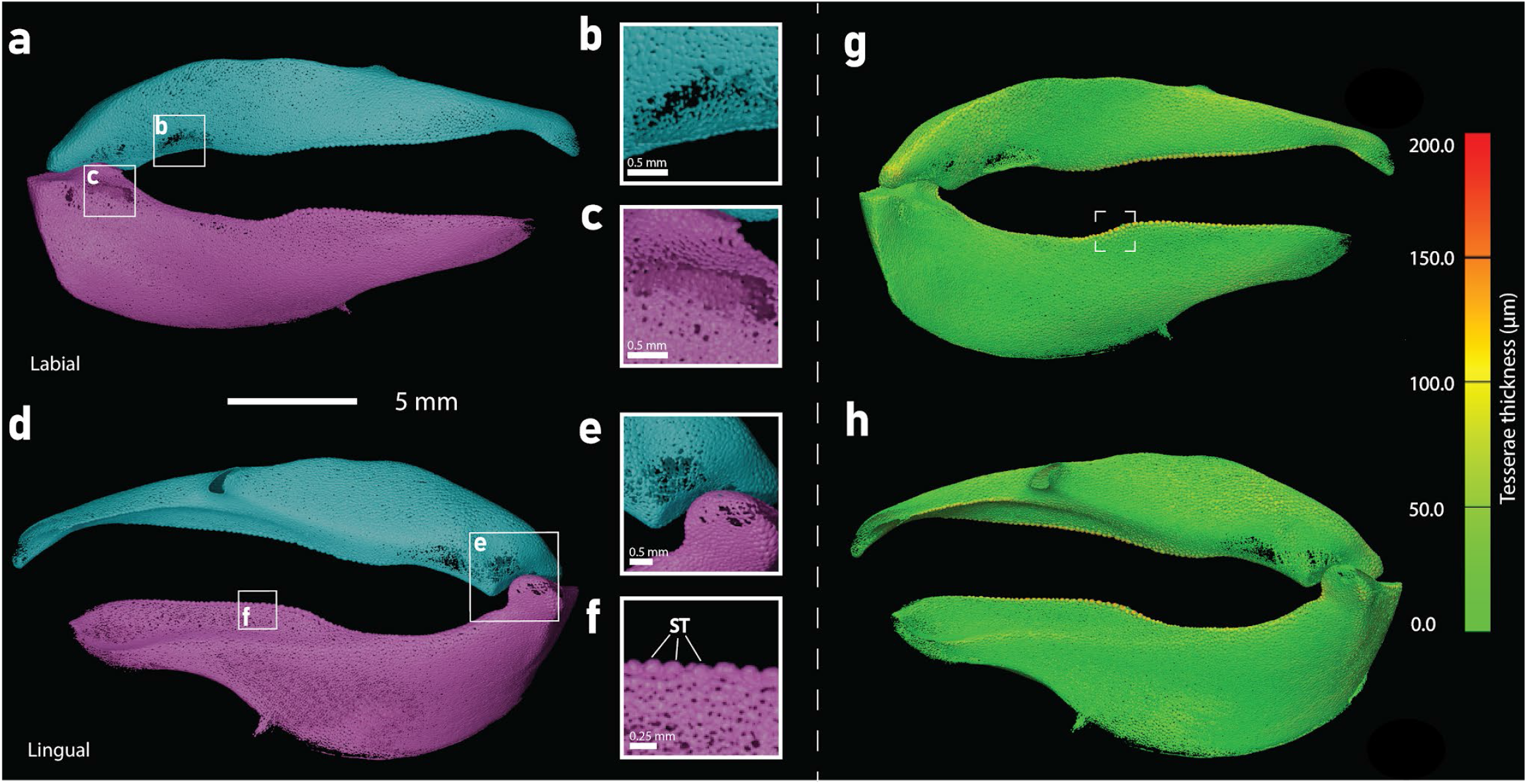
3D render of *Scyliorhinus canicula* jaws. The upper jaw (turquoise) and lower jaw (magenta) are shown in labial **(a)** and lingual **(d)** views. Magnified regions of interest **(b, c, e, f)** highlight Specialised Tesserae (ST) and zones of lower mineralisation. Spatial thickness map of the TC is shown in labial (**g**) and lingual (**h**) view with the colour scale saturating at maximum thickness. The white box on panel (**g**) is where the thickest region is located.

### Squalomorphs and holocephalans

#### Squatina oculata

There is a high degree of heterogeneity in tesserae shape, size and mineralisation across both the upper and lower jaws (Figure 15a, d). For the majority of the tessellated cartilage, tesseral pores and spokes are absent, and thus appear fused together (Figures 14a and 15a, d, f). Towards the central regions of the jaws, tesserae appear polygonal with small tesseral pores and narrow spokes, with an average max width on the upper jaw of 286 µm and lower jaw of 253 µm, giving an overall average of 270 µm, making them the largest tesserae in this study by almost 100 µm (Table 4;Figure 14b and Figure 15a, d). The tesseral layer is generally complete, with the presence of a gap at the orbital process of the upper jaw (Figure 15d, e). The average thickness of the layer is 63 µm with a maximum of 260 µm located at the proximal end of the upper jaw. This is the largest average and maximum tesseral layer thickness of any taxa in this study (Table 4; Figure 15g, h). There are numerous large canal pores present on both the upper and lower jaw, with the largest appearing proximally on the lower jaw with a max width of 550 µm, equalling the largest canal pore of the study in *Ginglymostoma* (Table 4; Figure 15a, b, d, f). These pores are associated with internal mineralisation (Figure 15e), which are likely rudimentary trabeculae that vary in development but none fully extend between two tesseral layers. No specialised tesserae were identified.

**Figure 14.**
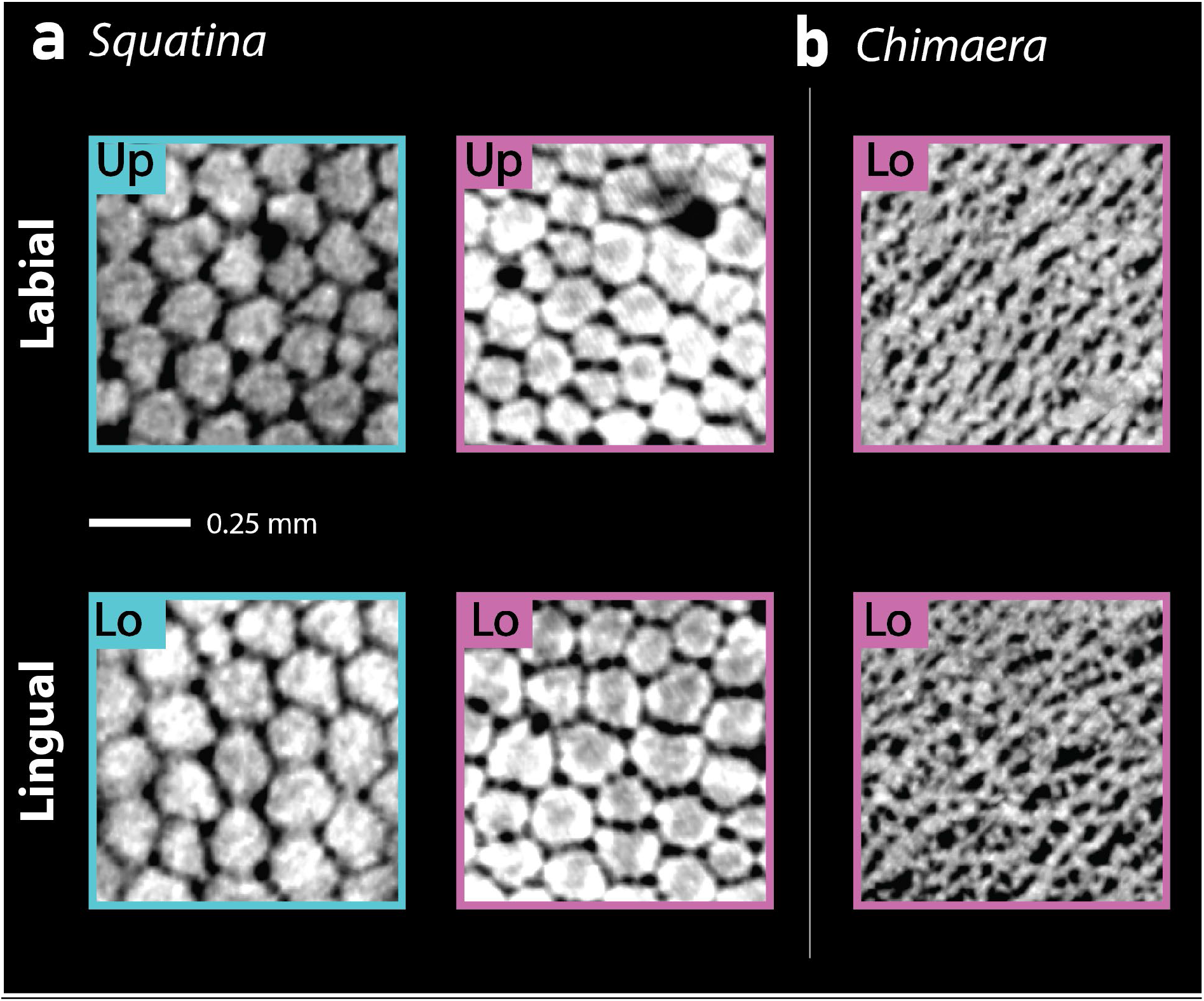
Virtual thick slabs of tesserae from the Upper (Up) and Lower (Lo) jaws of the one squalomorph and one holocephalan taxa in this study **(a)** *Squatina* and **(b)** *Chimaera* in labial and lingual view.

**Figure 15.**
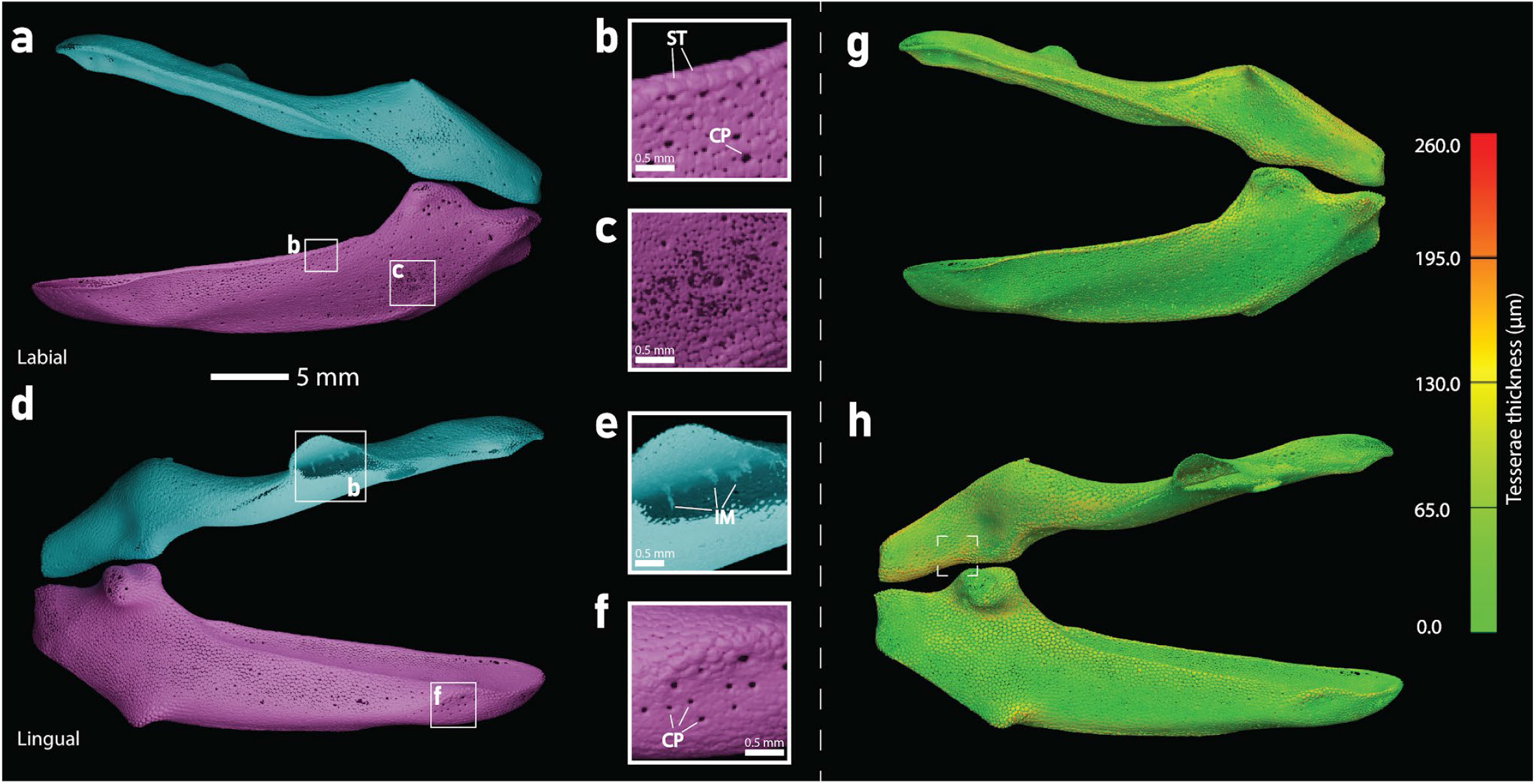
3D render of *Squatina oculata* jaws. The upper jaw (turquoise) and lower jaw (magenta) are shown in labial (**a**) and lingual (**d**) views. Magnified regions of interest **(b, c, e, f)** highlight Canal Pores (CP), internal mineralisation (IM) and zones of lower mineralisation. Spatial thickness map of the TC is shown in labial (**g**) and lingual (**h**) view with the colour scale saturating at maximum thickness. The white box on panel (**h**) is where the thickest region is located.

#### Chimaera monstrosa

Only the Meckel’s cartilage is included in this study as the palatoquadrate is fused with the neurocranium (Didier, 1995) and cannot be segmented out in the scans. There are no polygonal tesserae present, instead tesserae tiles are reduced with the spokes becoming dominant (stellate), this is best seen in the thick slabs (Figure 14b; Figure 16). The tessellated cartilage varies in appearance throughout the jaw, proximally the tessellated cartilage appears stellate and aborally it appears fused and like a thin sheet with small intermittent tesseral pores (Figure 14b and Figure 16a-f). The tesseral layer is incomplete in many regions, the most notable being the oral margin (Figure 16 a, d). The average thickness of the layer is 17 µm with a maximum of 80 µm located centrally on the labial surface (Table 4; Figure 16g, h). There are no canal pores present, no mineralised structures seen internally and no specialised tesserae.

**Figure 16.**
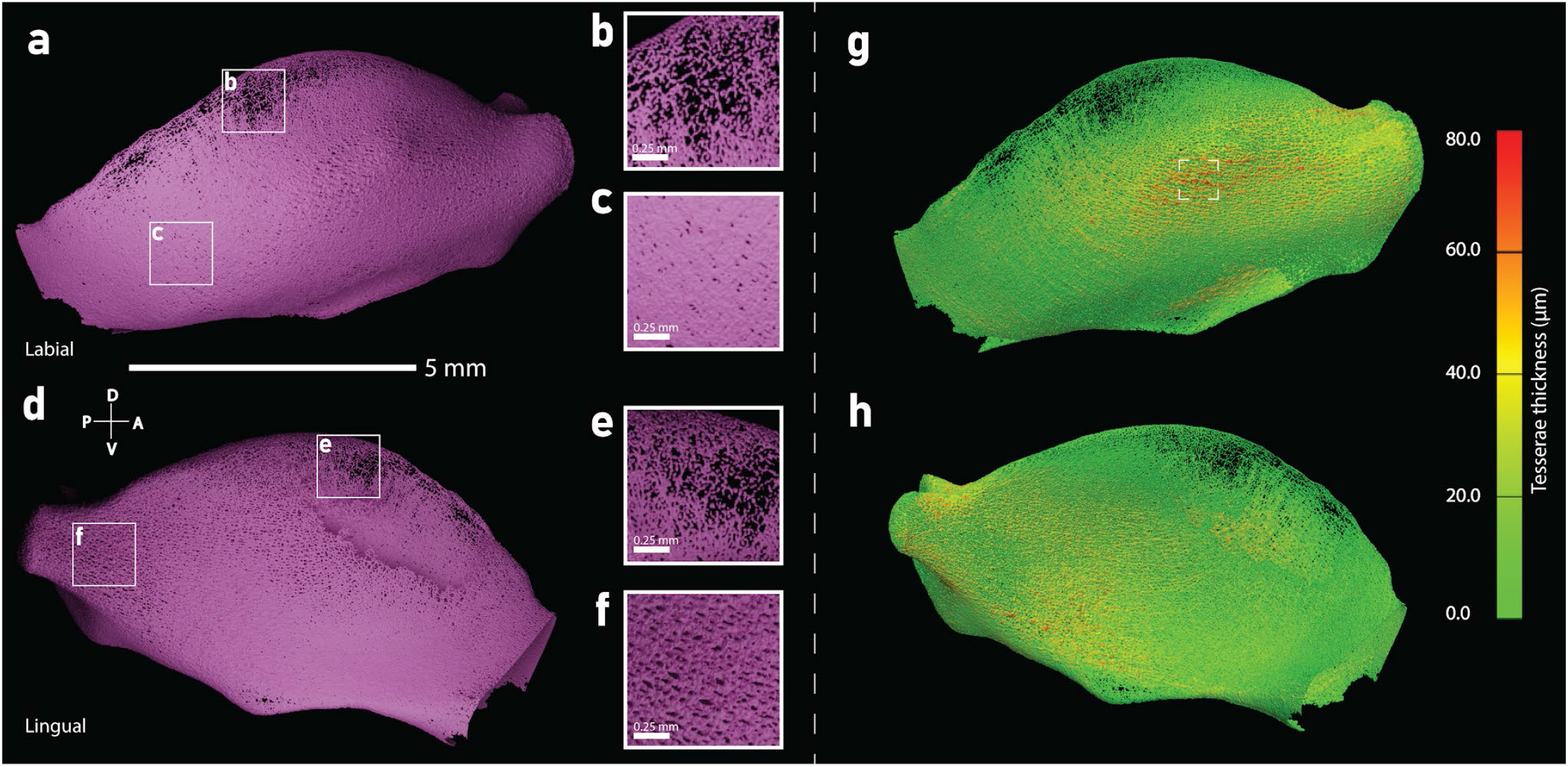
3D render of *Chimaera monstrosa* lower jaw (magenta) is shown in labial **(a)** and lingual **(d)** views. Magnified regions of interest **(b, c, e, f)** highlight tesserae and zones of lower mineralisation. Spatial thickness map of the TC is shown in labial (**g**) and lingual (**h**) view with the colour scale saturating at maximum thickness. The white box on panel (**g**) is where the thickest region is located.

## Discussion

In agreement with previous studies (Berio et al., 2021; Clark et al., 2022; Dean et al., 2009; Seidel et al., 2020a, 2016), we have shown that tesserae are typically tightly abutting with no gaps, except in localised regions of patchy tessellation (Figures 4,7,9,11-13,15); this means that larger jaws should have larger and/or more tesserae relative to smaller ones to account for the difference in surface area. Recent work on the stingray *U*. *halleri (Seidel et al., 2016)* has shown that during development, the number of tesserae on a given piece of the skeleton is relatively constant, with tesserae mostly increasing in size not number to accommodate the growing skeleton. In contrast, in our comparisons of batoid species with different sized jaws, tesserae were well-developed, polygonal, and relatively homogeneous in size and shape across both upper and lower jaws (Figures 3-7). Notably, these traits were observed in *Torpedo torpedo* despite its much slighter jaw morphology and distinct feeding habits relative to the other three taxa (Figures 4-7). Tesserae in the closely-related *Torpedo californica* have been described as irregular in shape (Seidel et al., 2016) which is in contrast to our findings for *T. torpedo*. Among the four batoids studied, tesserae were largest in *T. torpedo* (mean width: 174 µm), which, given previous demonstrations of tesserae enlargement during ontogeny (Clark et al., 2022; Yang et al., 2024), likely reflects its slightly later ontogenetic stage (Table 2 and 3). Interestingly, *Zanobatus schoenleinii* possessed the largest tesserae relative to jaw length of any taxon in this study (Table 4; SM Figure 11). Because its tesserae have not been previously described, the underlying developmental drivers remain unclear, highlighting the need to examine older ontogenetic stages.

Tesseral layer thickness also varied among the batoids; *Z. schoenleinii* and *Potamotrygon orbignyi* exhibited tesseral layers less than half the thickness of most taxa in this study (mean: 17 µm and 18 µm respectively (Table 4; SM Figure 13). Small trabeculae were present in the lower jaws of *Rhinobatos cemiculus* and *P. orbignyi* (Figures 4c, 5c), with rudimentary trabeculae in Z. *schoenleinii* and none in *T. torpedo*. Based on observations in mature batoids, additional trabeculae will likely develop later in ontogeny (Clark et al., 2022). This may not be the case with *T. torpedo*, trabeculae are strongly associated with canal pores and none are observed for the specimen in our study. Although multi-layered tesserae have been reported in older batoid taxa (Clark et al., 2022; Dean et al., 2009; Summers, 2000), no multi-layering was observed here, further supporting the idea that multi-layering develops later in ontogeny.

Among galeomorph sharks, *Ginglymostoma cirratum* was particularly notable, exhibiting the highest degree of tesseral morphological heterogeneity among all taxa studied (Figures 3, 8, 9, 14). In some regions, tesserae were polygonal with thin spokes; in other areas, the central disc was reduced, appearing stellate with spokes dominating (Figure 8a); in still other regions, tesserae appeared fused. The average tesseral layer thickness was 49 µm, among the highest in this study (Table 4). Most unexpectedly, an extensive array of trabeculae was present in both jaws, including a large, branching trabecular cluster in the lower jaw (Figure 10). To our knowledge, trabeculae have never been described in shark jaws, yet these trabeculae were the most developed and numerous of all specimens in this study. The reasons for such intricate trabeculae are unclear, yet we hypothesise that their presence and morphology relate to the unique mechanical demands of nurse shark feeding ecology. Trabeculae have typically been associated with durophagy, arranged like pillars buttressing the jaw cortex beneath the teeth in species that eat hard foods (Clark et al., 2022; Dean et al., 2006; Summers, 2000). In contrast *G. cirratum* is a suction feeder, able to generate impressive subambient pressures rapidly enough to induce cavitation, with the fastest mandibular depression recorded for any shark (Motta et al., 2002). These behaviours suggest that jaws experience lateral buckling stresses rather than occlusal stresses: indeed, the trabeculae we observed are oriented labio-lingually, rather than occlusally. Similar “truss trabeculae” have only been documented in the jaws of the electric ray, *Narcine bancrofti* (Dean et al., 2006), which deploys its jaws like a long post-hole-digger to excavate worms (Dean & Motta, 2004). The fact that this cross-bracing morphology for trabeculae is now demonstrated to be common between two distantly-related chondrichthyans suggests this more subtle role —requiring less robust trabeculae than durophagy— may be more widespread than appreciated. The interindividual variability in feeding kinematics seen in *G. cirratum* (Motta et al., 2002) also begs the question whether interindividual variations in tesserae and the trabeculae will be observed. Further examination of specimens across ontogeny —ideally under controlled feeding regimes— is needed to elucidate further on the role and plasticity of trabeculae in *G. cirratum,* in addition to the need to examine surrounding skeletal elements, like the labial cartilages and hyoid arch which also play crucial roles during suction, to see if trabeculae occur there also. In addition, broader sampling across Orectolobiformes is needed to determine if this trait extends beyond *G. cirratum*.

We observed substantial heterogeneity in tessera size, shape, and mineralisation among the carcharhiniform sharks studied (Figures 8, 11-13). *Mustelus mustelus*, for example, exhibited no discernible tesseral discs and contained unmineralised patches. For *M. mustelus*, this limited TC development is surprising given that its early ontogenetic diet is dominated by crustaceans (Compagno, 1990). This poor development stands in stark contrast to other durophagous elasmobranchs in our study (Figures 4, 5, 7, 12). *Mustelus mustelus* even possesses molariform teeth specialised for crushing hard prey (Compagno, 1984), making this developmental lag particularly puzzling. The presence of multi-layered TC in an older individual (H. Byrne, pers. obs.) further highlights the substantial ontogenetic changes in the taxon. Consequently, examining specimens at similar and later ontogenetic stages as well as other species within the genus *Mustelus* will be essential to better understand how jaw morphology changes with growth, and potentially environment and feeding. Conversely, *Rhizoprionodon acutus* possessed well-developed tesserae with thick spokes alongside specialised tesserae*. R. acutus* undergoes a reverse dietary shift from teleosts to crustaceans later in life; investigating older specimens will clarify whether trabeculae and multi-layered TC develop to accommodate increased durophagy (Ba et al., 2013; Sen et al., 2018). *Scyliorhinus canicula* occupied an intermediate state, featuring small, poorly developed tesserae (but with specialised tesserae) and recorded the lowest TC thickness among the selachians at 33 µm (Table 4; Figures 8,11-13, SM Figure 12-13). This result for *S. canicula* aligns with previous work demonstrating slower rates and lower overall levels of mineralisation in adults of this species compared to batoids (Debiais-Thibaud, 2019). What can we say about the specialised tesserae?

While Squatiniformes are characterised by relatively large, highly mineralised tesserae, these traits have been documented primarily in fossil specimens (Cabrera et al., 2012; Maisey et al., 2020b), leaving extant forms understudied (Berio et al., 2021). Our observations generally align with the findings from the aforementioned studies (Figures 15b, 16). The tesserae of *Squatina oculata* were the largest of any of the taxa in the study (273 µm), and both jaws contained extensive areas of fused tesserae indicative of high mineralisation (Table 4; Figure 15b,16). Tesserae size relative to jaw length is low, but is the highest relative to body length (Table 4; SM Figures 11-12). Tesseral layer thickness was the highest in our study (63 µm), and is high relative to tessera size (Table 4; SM Figure 13). The reason for the large and highly mineralised tesserae in Squatiniforms remains unclear, but it may relate to the mechanical stresses generated during feeding. Despite substantial research on Squatiniformes, largely driven by the critically endangered status of many taxa, only a single study has documented their ambush predation. Fouts and Nelson (1999) observed rapid ambush strikes lasting just 30–100 ms, accompanied by cranial elevation reaching up to 90°. These explosive movements likely induce substantial mechanical stress, high mineralisation and enlarged tesserae may therefore provide increased support and stiffness to withstand this stress. Given that both batoids and squatiniforms possess well-mineralised tesserae alongside dorsoventrally flattened body plans, assessing whether highly mineralised tesseral layers occur in other dorsoventrally flattened sharks (e.g., *Orectolobus, Pristiophoru*s) is of great interest.

The tessellated cartilage of the holocephalan *Chimaera* differed greatly from that of the other taxa in our study, with no polygonal tesserae and large gaps in the tesseral layer (Figures 3, 8, 14). These differences have been noted in several studies, with debate surrounding whether tesserae are even present in holocephalans (Atake et al., 2025; Clarac et al., 2025; Seidel et al., 2020a). Other studies (Clarac et al., 2025; Seidel et al., 2020a) have noted chimaeroid tesserae to be small and stellate, which may explain why our scans do not resolve them; a future study of holocephalan skeletal tissue at sub-micron resolution will help settle this debate. Similar poorly developed tesserae have been documented in the holocephalan *Hydrolagus colliei* and *Callorhinchus milii* (Berio et al., 2021; Pears et al., 2020), but also in two deep-water squalomorphs (*Etmopterus spinax* and *Centroselachus crepidater;* Berio et al., 2021), along with a near absence of tessellation in some parts of the skeleton of large coldwater sharks (e.g., *Somniosus*, *Hexanchus*, *Notorynchus*; Dean et al., 2015). These collected observations underline the need for further investigation into potential environmental drivers of tesseral layer development, particularly in the face of our changing oceans.

Specialised tesserae were identified in both the batoid and galeomorph taxa examined in this study (*Zanobatus*, *Rhizoprionodon*, and *Scyliorhinus*). These structures were consistently larger than typical tesserae and were notably wider than they were tall (Table 4). This morphology contrasts with the voussoir and columnar tesserae of mature specimens, which are traditionally characterised as being taller than they are wide (Clark et al., 2022; Maisey et al., 2020a). Although the specialised tesserae in our study were located in regions where voussoir and columnar forms are expected, examining these taxa across an ontogenetic series is necessary to conclusively link them. Additionally, while voussoir tesserae have been previously described in Squatiniformes, they were not definitively identified in our *Squatina* specimen (Maisey et al., 2020a). Some tesserae exhibited borderline morphology; however, given the early developmental stage of our specimen, it is highly possible that these structures fully differentiate and become identifiable later in ontogeny.

## Conclusions

The tesseral layer at early stages of ontogeny varies greatly across the four major chondrichthyan groups studied. Among the batoids, tesserae are well-developed and defined early in ontogeny, regardless of feeding habits and jaw morphology, echoing the findings of previous studies (Dean et al., 2009; Debiais-Thibaud, 2019). The unified characteristics could be due to similar feeding habits and/or body morphologies. There are no unifying traits among the TC of the galeomorphs, with an unexpected intricate internal trabeculae array found in *G. cirratum* likely related to the high pressures generated as a specialised inertial suction feeder. This is the first time trabeculae have been described outside of the batoids. The differences between the galeomorphs studied could be due to differences in feeding habits, this would need to be explored further using older individuals.

Although no novel TC traits were observed in this study, specialised tesserae were recorded in both batoids and galeomorphs. These specialised tesserae also appear to exhibit a reversed aspect ratio compared to those described in previous studies (voussoir and columnar), that is, they are wider than they are tall. An increase in height may therefore develop later through ontogeny (Clark et al., 2022; Maisey et al., 2020a). The ontogenetic development of trabeculae appears to vary, with only one or two developed in neonate/juvenile batoids compared to the extensive array seen in *G. cirratum* at the same early ontogenetic stage. No multiple layers of TC were observed in any of the specimens, strongly suggesting that this is a feature developed later in ontogeny.

In light of this broader sampling, generalised models of TC development based on single groups or limited taxa fail to capture the true diversity of TC morphology. This study establishes a foundation for future research encompassing additional taxa and ontogenetic stages to evaluate TC morphology more thoroughly and assess other drivers, including habitat, body size, and sex. Furthermore, it provides a framework to investigate more difficult-to-interpret tesserae in fossil sharks and ultimately examine the evolution of TC across 430 million years of chondrichthyan history.

## Acknowledgements

We would like to sincerely thank Esther Dondorp (Curator for fishes at Naturalis Biodiversity Center) for helping provide access and permission to scan the specimens in this study.

## Data Accessibility

All micro-CT scans used in this study are publicly available on MorphoSource (https://www.morphosource.org/projects/000929153?locale=en), and our datasets and R scripts are hosted on GitHub (https://github.com/hbyrne628/TC-scans.git).

