## Supplementary Figures 1-14 for "Early Ontogenetic Development of Tessellated Cartilage in Chondrichthyans"

**SM figures**
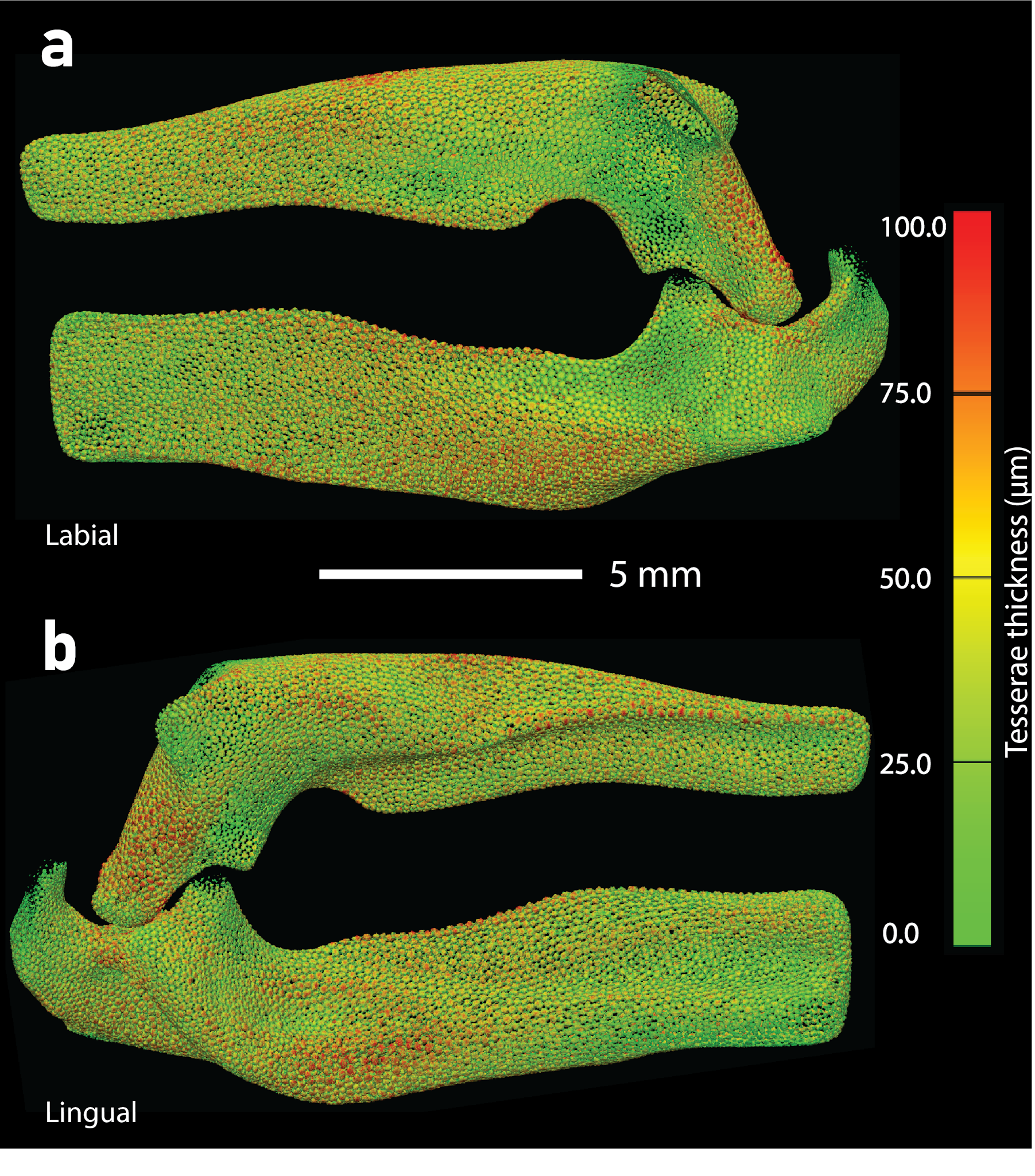

**SM Figures 1 and 2** Spatial thickness map of the tessellated cartilage is shown for the jaws of *Glaucostegus* (left) and *Potamotrygon* (right). Color scale saturates at 100 µm (red) to emphasise relative thickness variations.
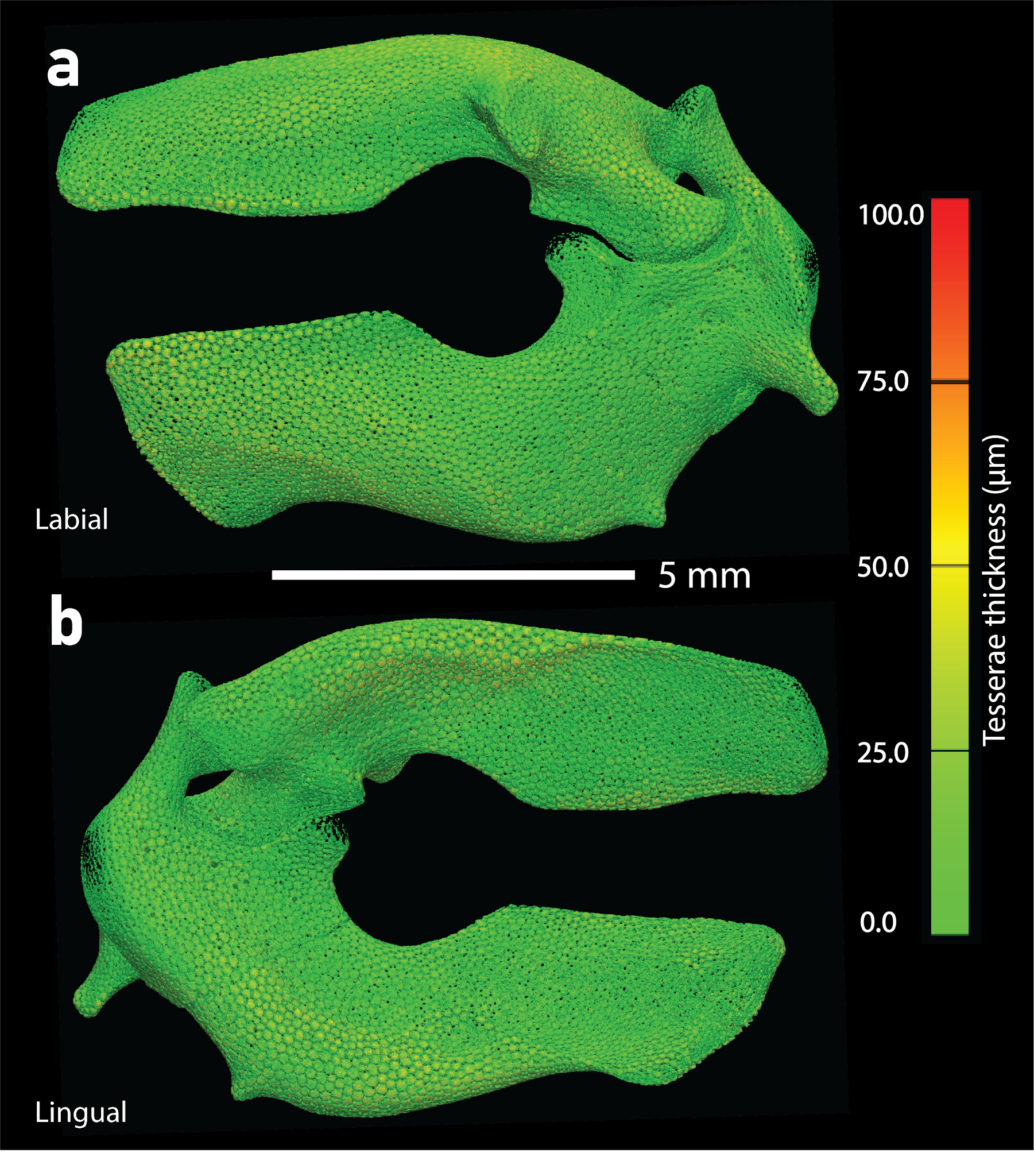

**SM Figures 3 and 4** Spatial thickness map of the tessellated cartilage is shown for the jaws of *Torpedo* (left) and *Zanobatus* (right). Color scale saturates at 100 µm (red) to emphasise relative thickness variations.
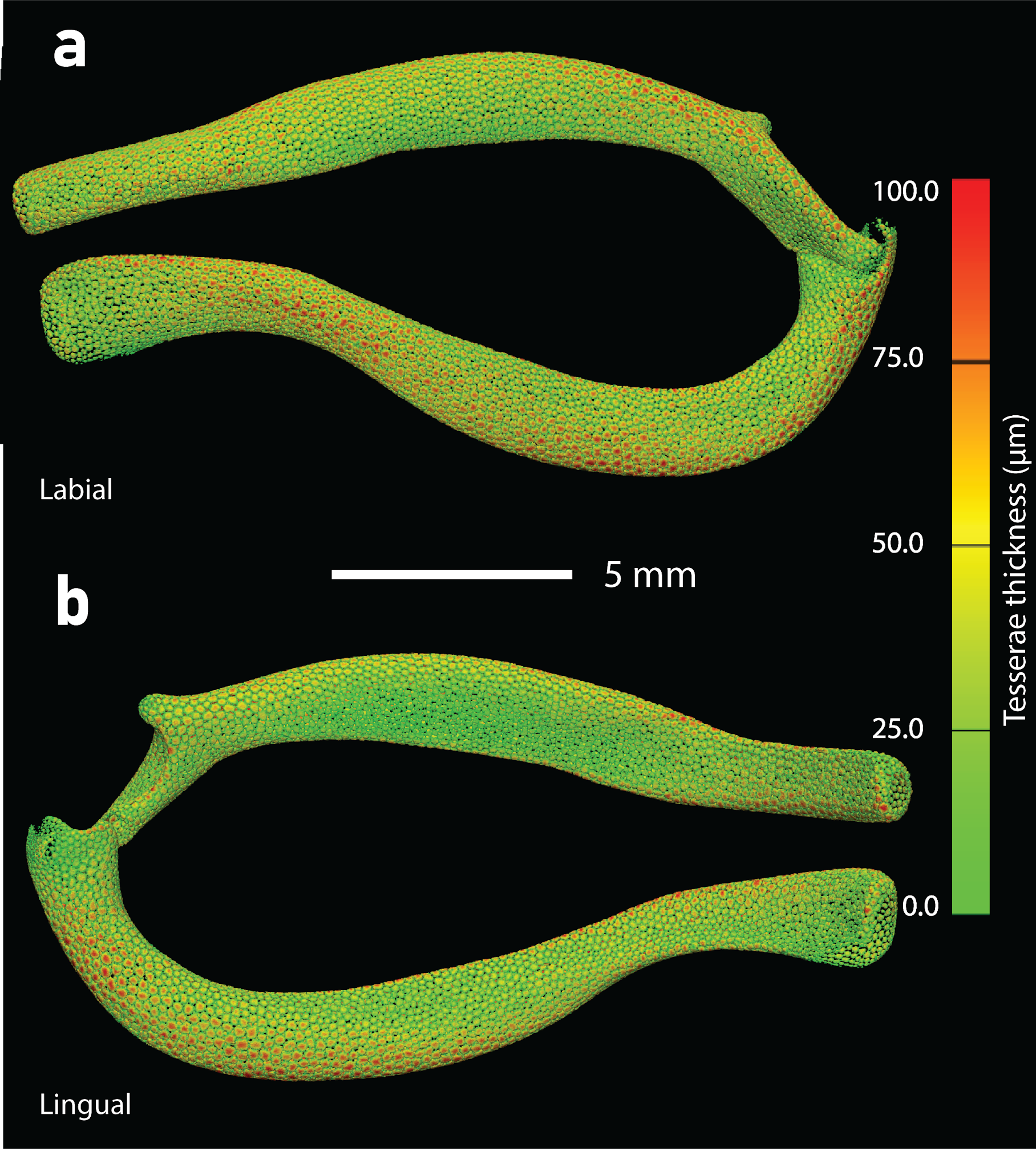

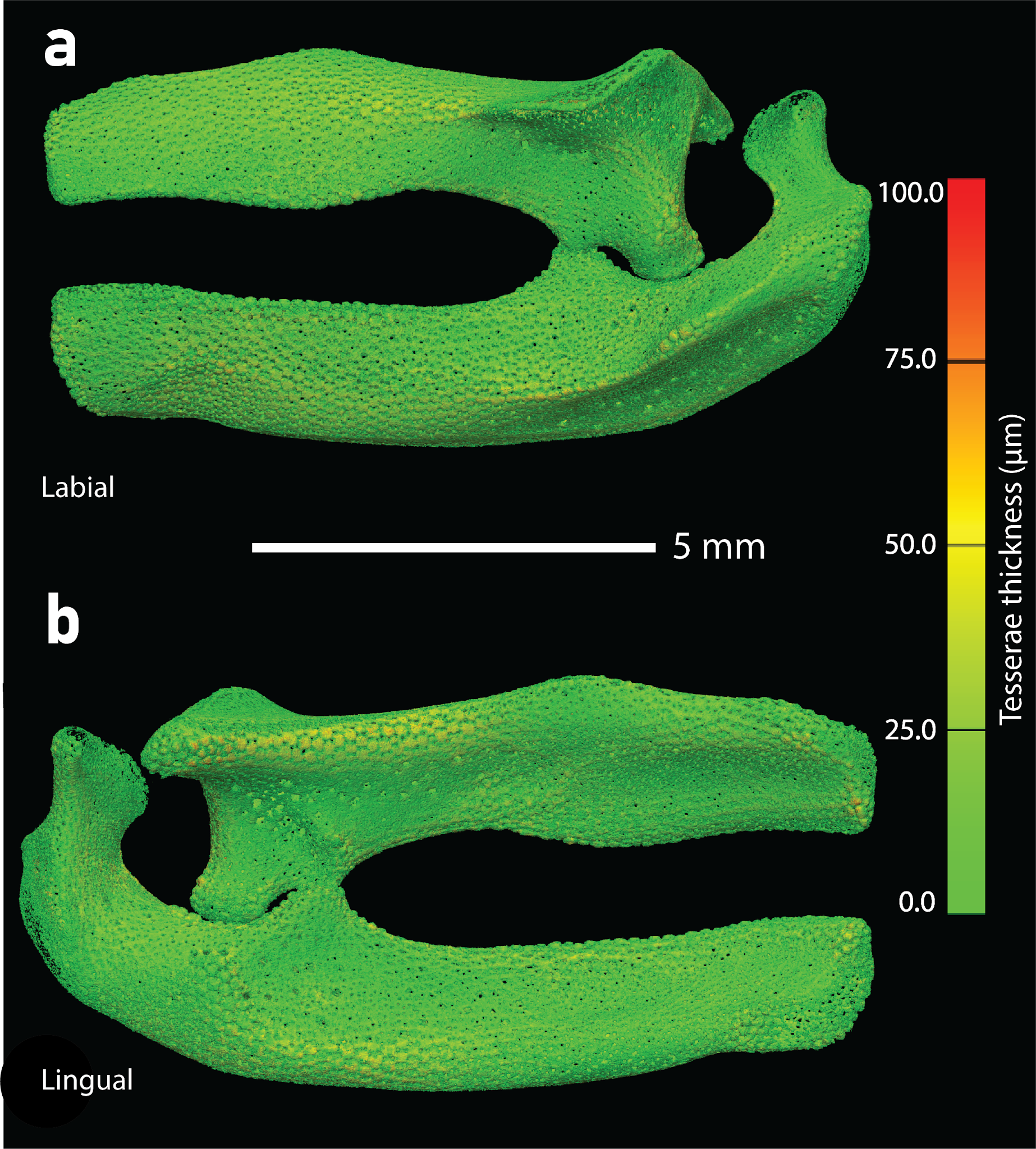

**SM Figures 5 and 6** Spatial thickness map of the tessellated cartilage is shown for the jaws of *Ginglymostoma* (left) and *Mustelus* (right). Color scale saturates at 100 µm (red) to emphasise relative thickness variations.
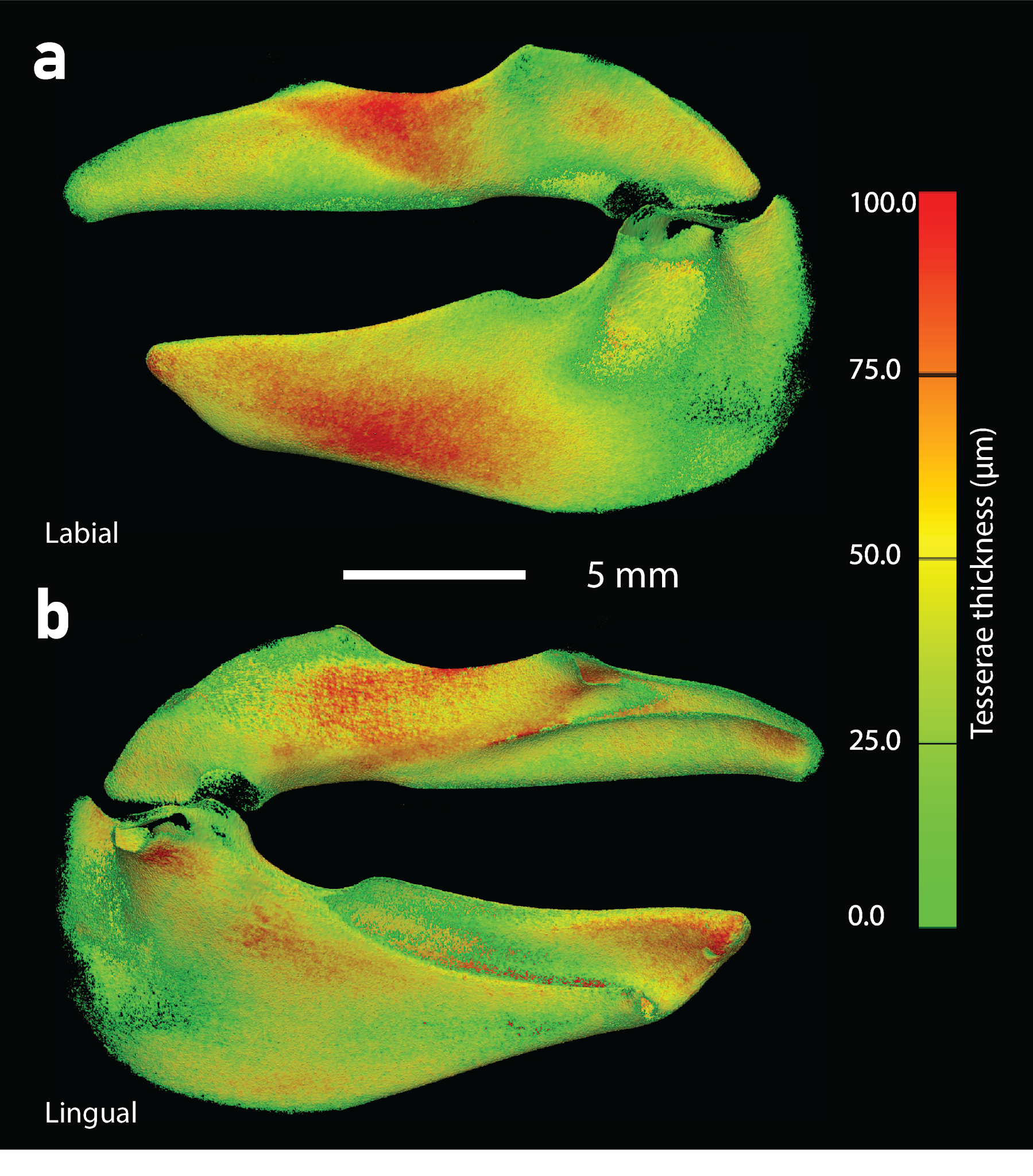

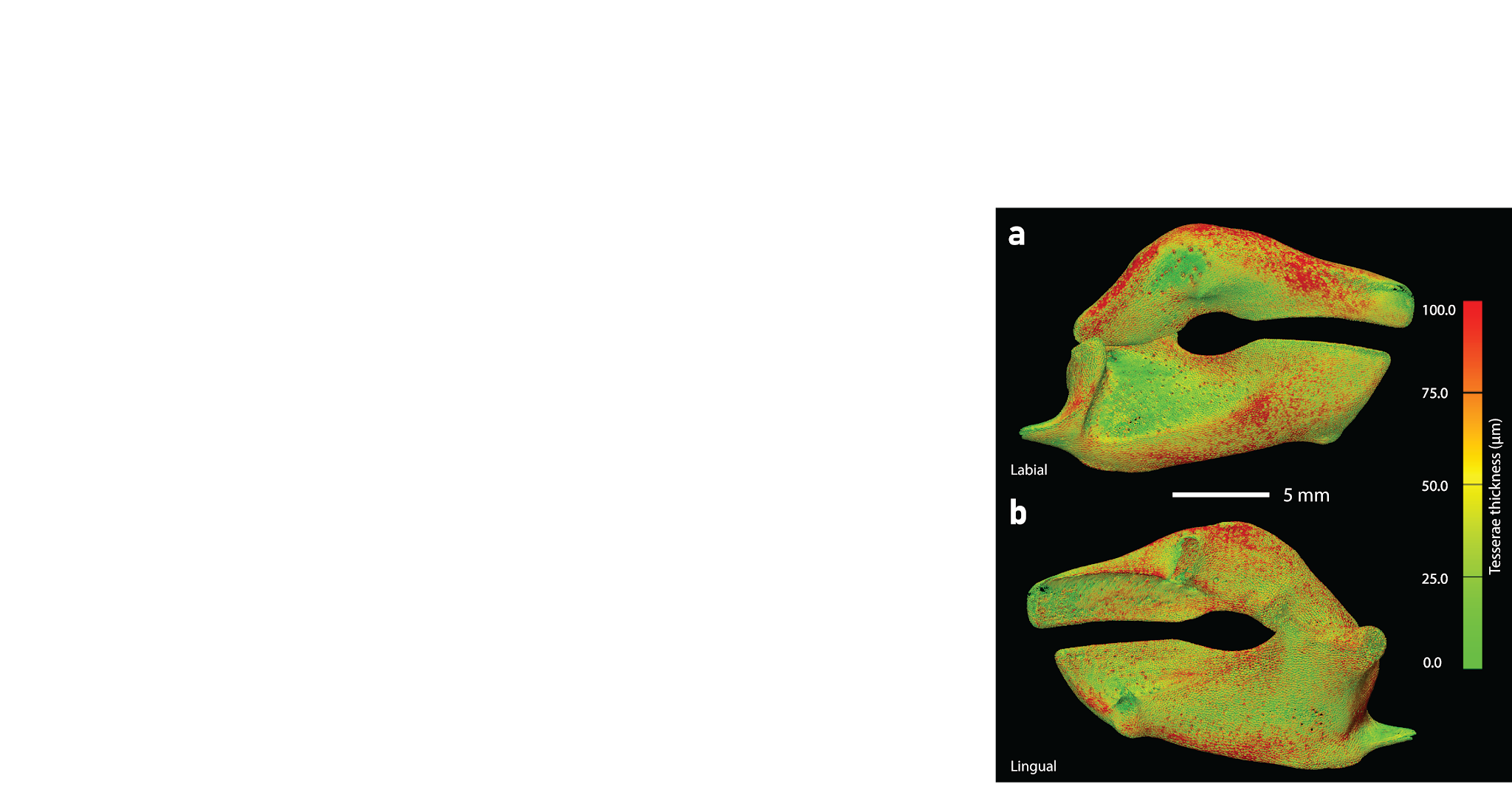

**SM Figures 7 and 8** Spatial thickness map of the tessellated cartilage is shown for the jaws of *Rhizoprionodon* (left) and *Scyliorhinus* (right). Color scale saturates at 100 µm (red) to emphasise relative thickness variations.
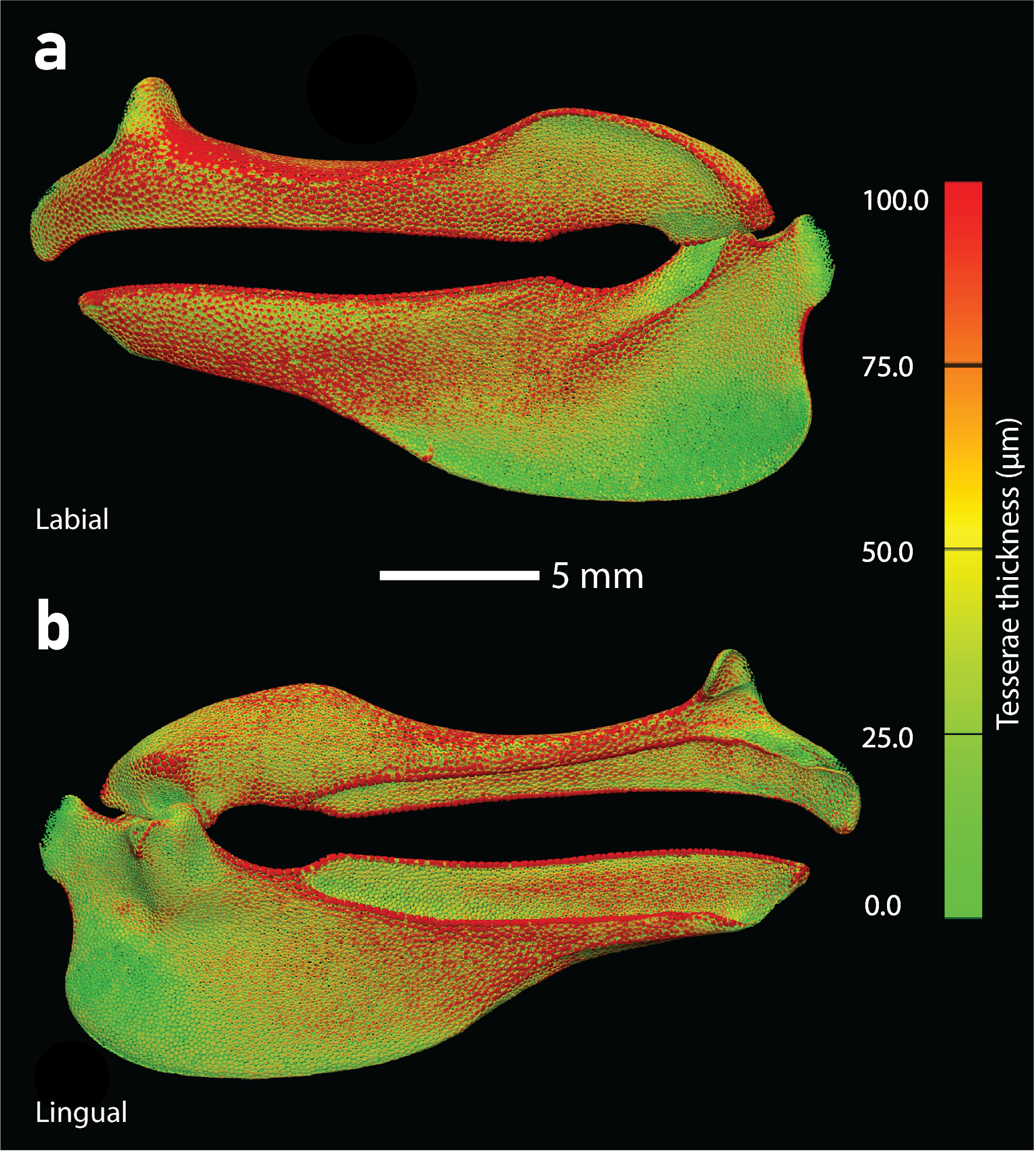

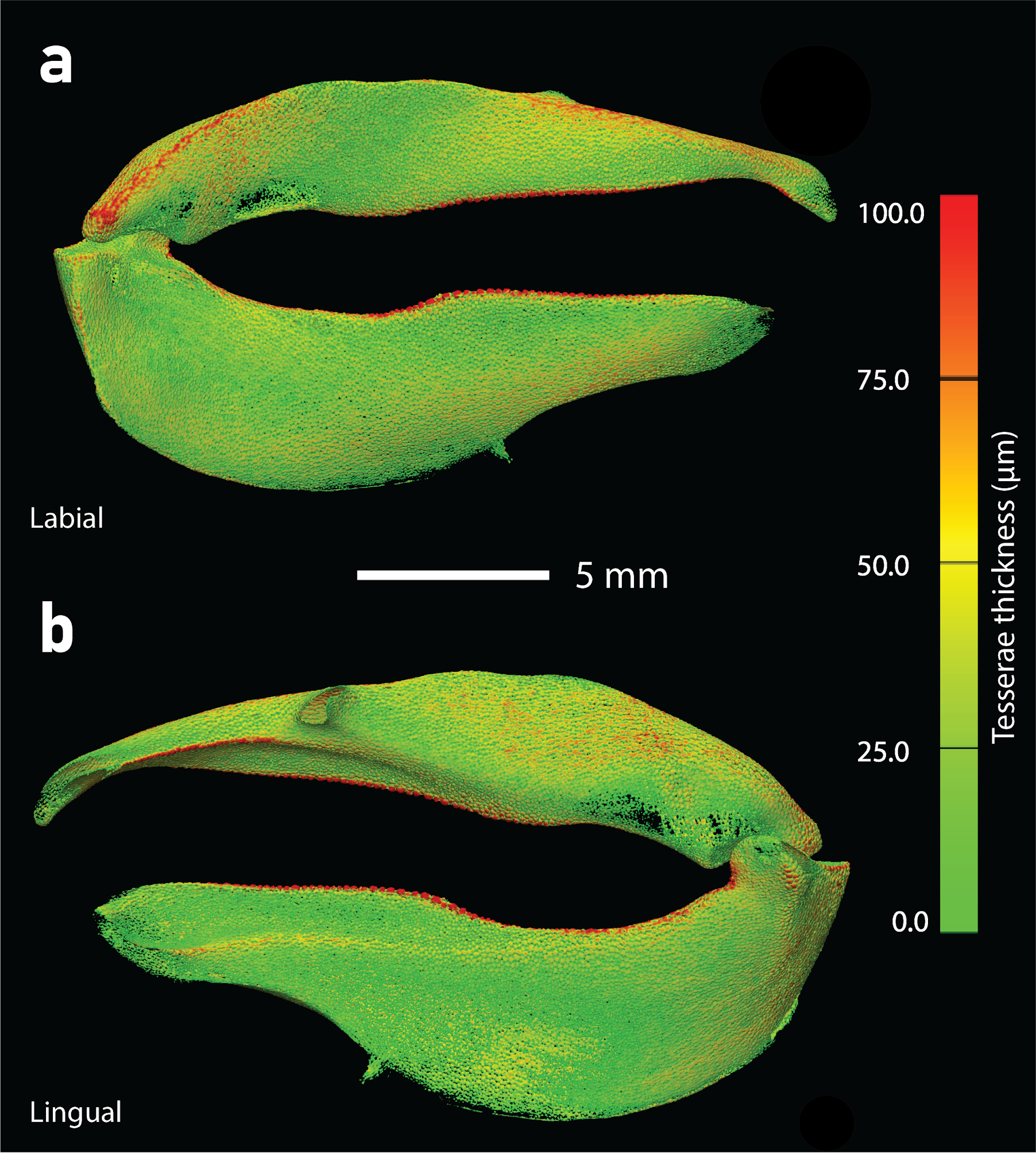

**SM Figures 9 and 10** Spatial thickness map of the tessellated cartilage is shown for the jaws of *Squatina* (left) and *Chimaera* (right). Color scale saturates at 100 µm (red) to emphasise relative thickness variations.
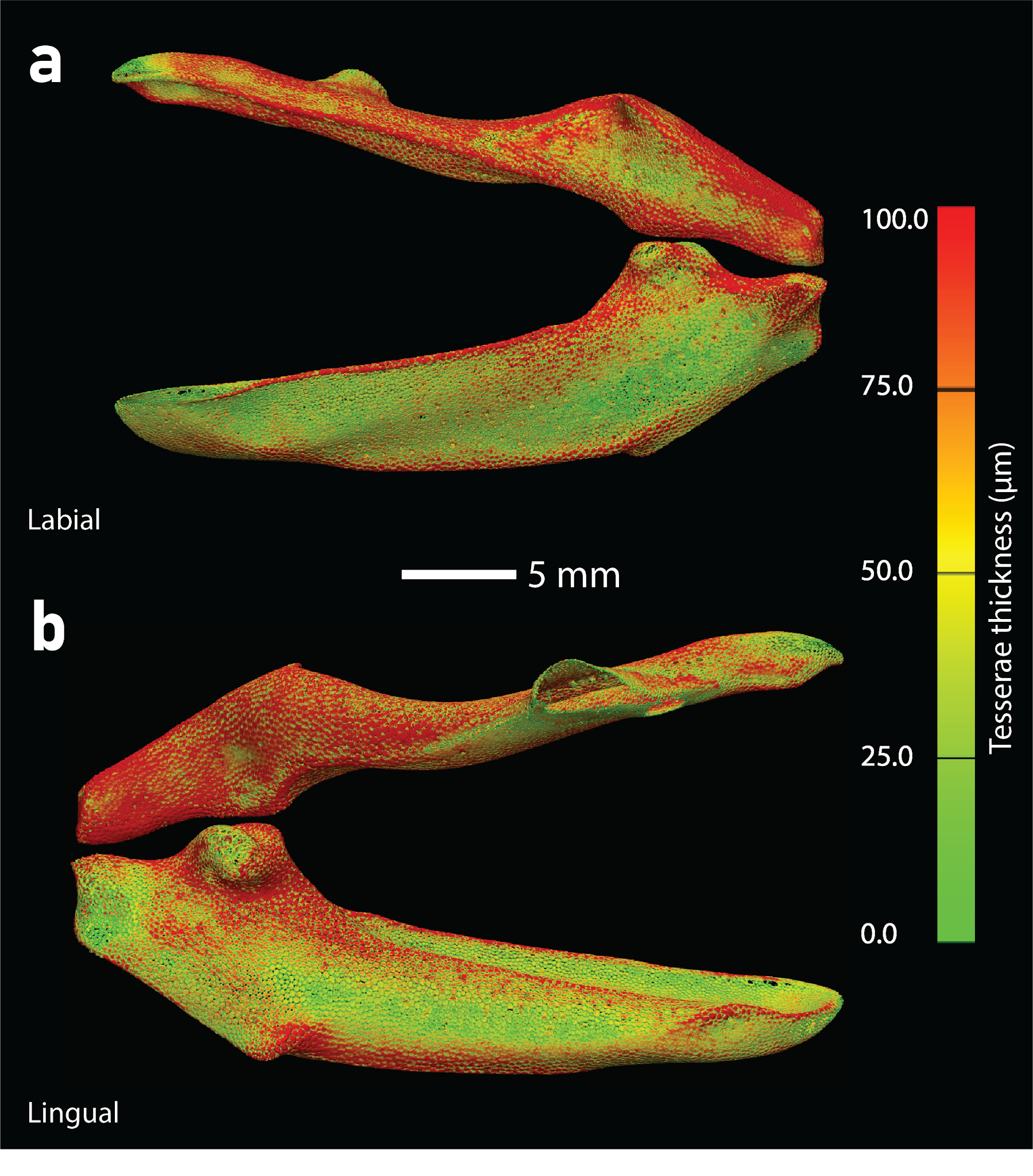

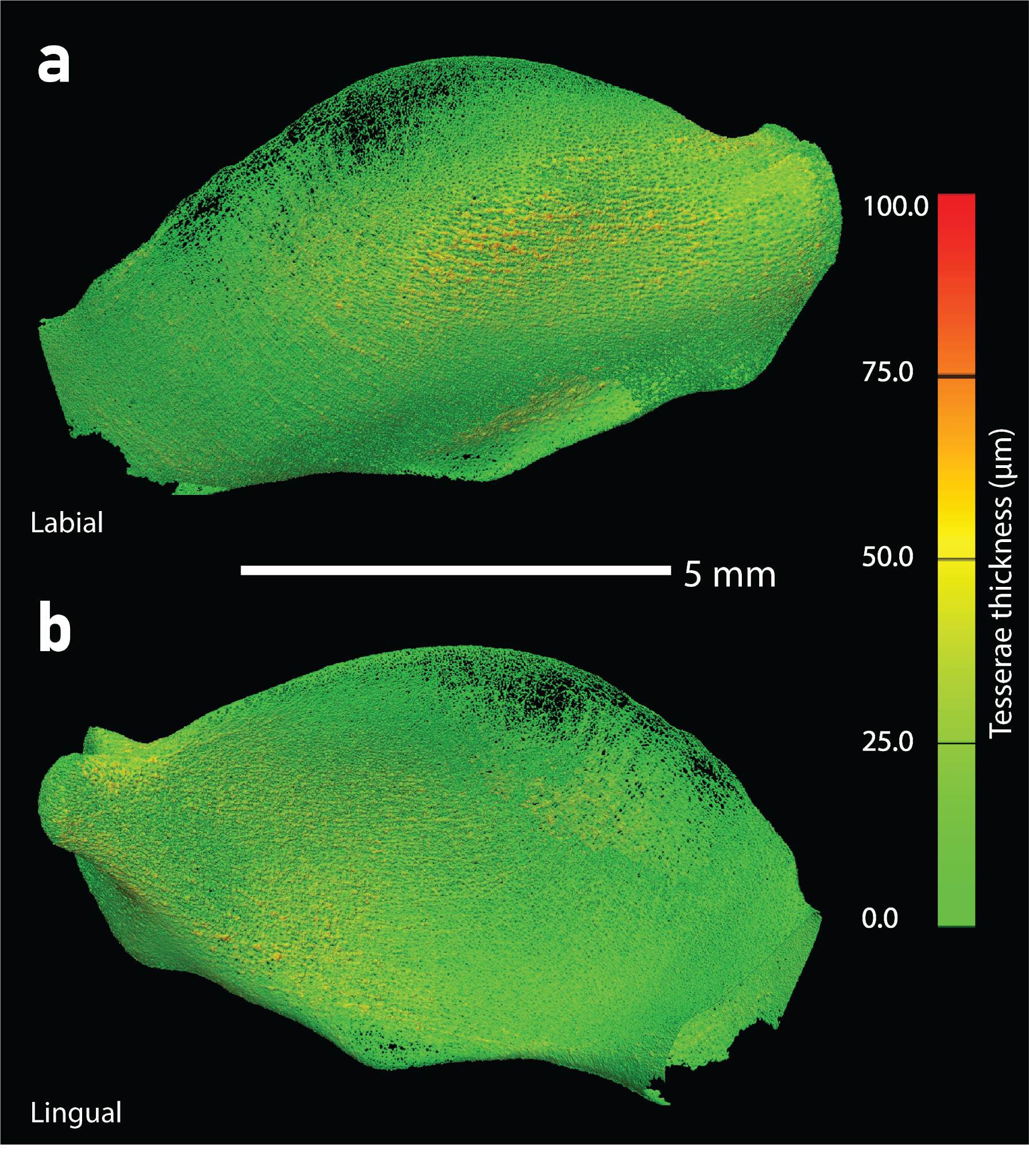

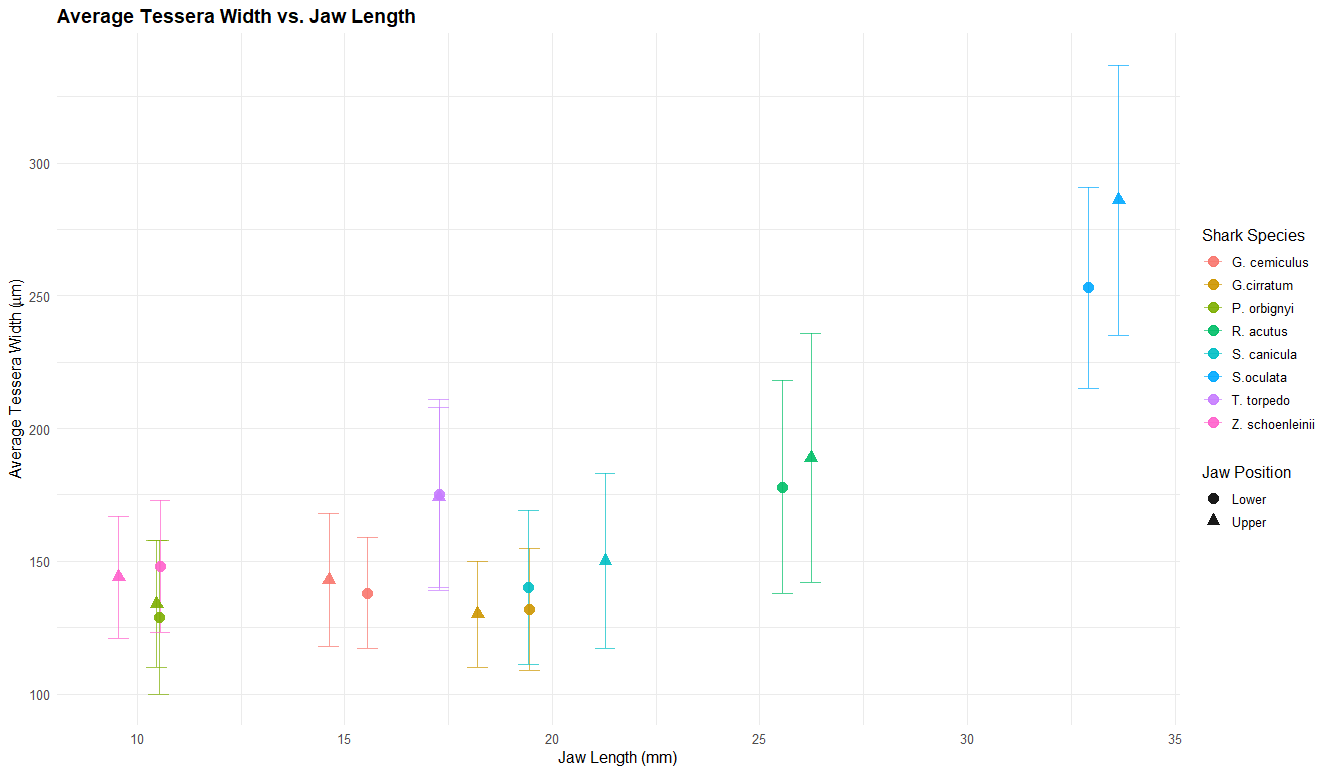

**SM Figure 11** Scatter plot with error bars displaying the average tessera width (y-axis) against jaw length (x-axis) for both the upper and lower jaw, data shown in Table 4.

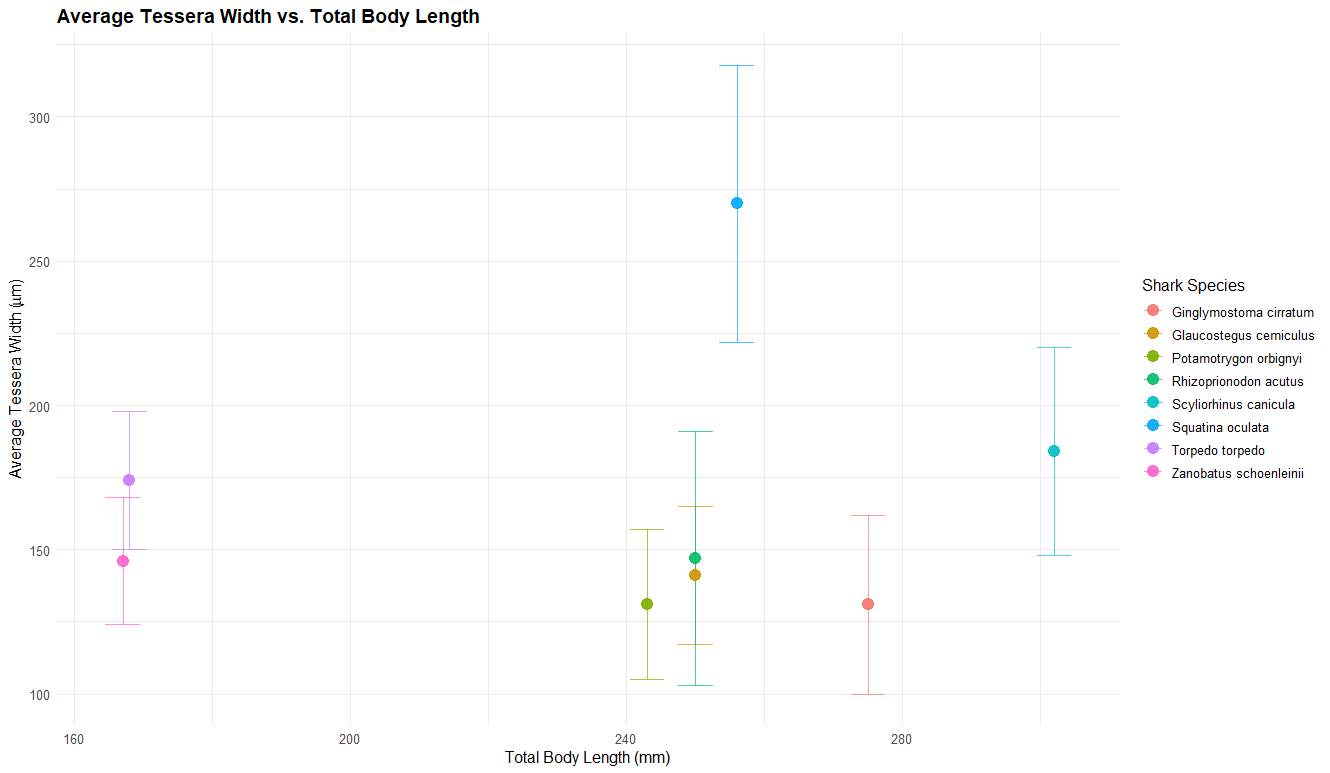

**SM Figure 12** Scatter plot with error bars displaying the average tessera width (y-axis) against total body (x-axis), data shown in Table 4.

**SM Figure 13** Scatter plot with error bars displaying the average tessera width (y-axis) against average tesseral thickness (x-axis), data shown in Table 4.
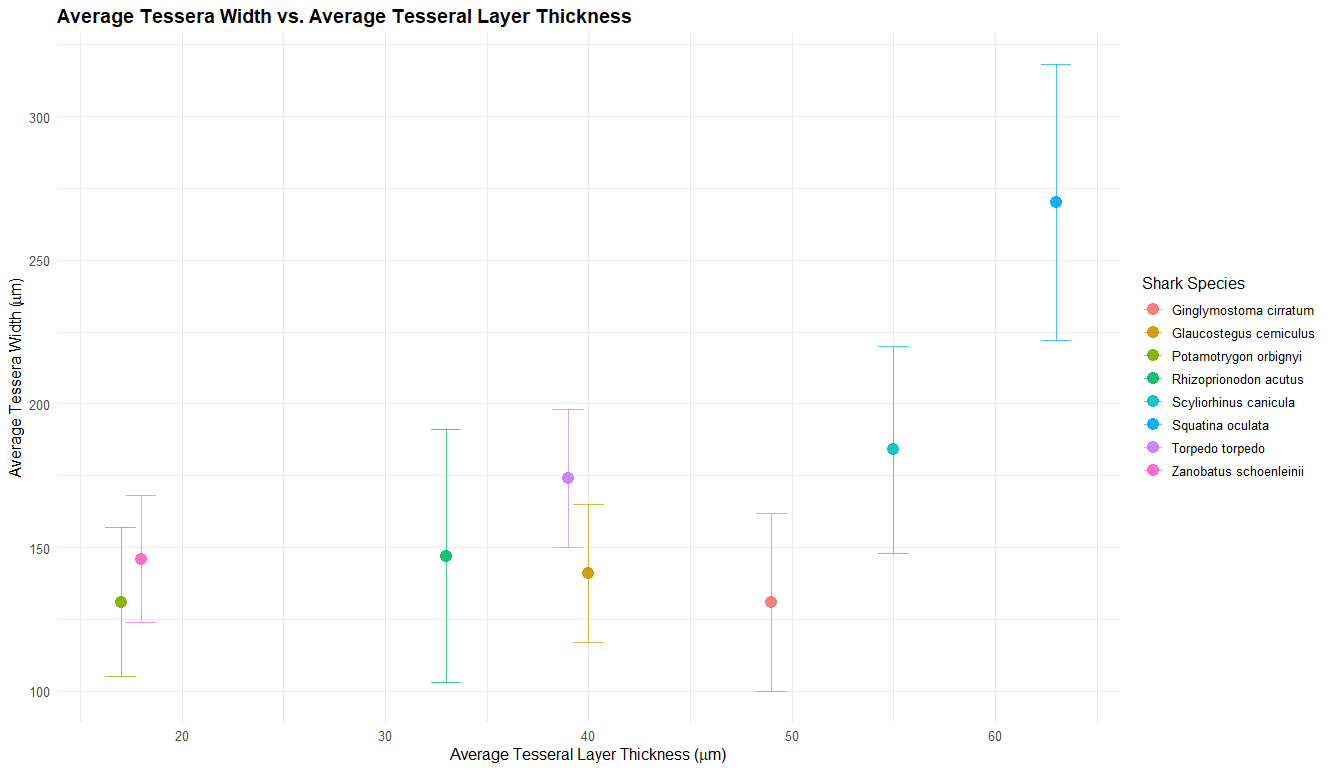
